# A SUMO-dependent regulatory switch connects the piRNA pathway to the heterochromatin machinery in *Drosophila*

**DOI:** 10.1101/2021.07.27.453956

**Authors:** Veselin I. Andreev, Changwei Yu, Juncheng Wang, Jakob Schnabl, Laszlo Tirian, Maja Gehre, Dominik Handler, Peter Duchek, Maria Novatchkova, Lisa Baumgartner, Katharina Meixner, Grzegorz Sienski, Dinshaw J. Patel, Julius Brennecke

## Abstract

Nuclear Argonaute proteins, guided by small RNAs, mediate sequence-specific heterochromatin formation. The molecular principles that link Argonaute-small RNA complexes to cellular heterochromatin effectors upon binding to nascent target RNAs are poorly understood. Here, we elucidate the mechanism by which the PIWI interacting RNA (piRNA) pathway connects to the heterochromatin machinery in *Drosophila*. Piwi-mediated stabilization of the corepressor complex SFiNX on chromatin leads to SUMOylation of its subunit Panoramix. SUMOylation, together with an amphipathic LxxLL motif in Panoramix’s intrinsically disordered repressor domain, are necessary and sufficient to recruit small ovary (Sov), a multi-zinc finger protein essential for general heterochromatin formation and viability. Structure-guided mutations that abrogate the Panoramix–Sov interaction or that prevent SUMOylation of Panoramix uncouple Sov from the piRNA pathway, resulting in viable but sterile flies in which Piwi-targeted transposons are derepressed. Thus, by coupling recruitment of a corepressor to nascent transcripts with its SUMOylation, Piwi engages the heterochromatin machinery specifically at transposon loci.

## INTRODUCTION

Heterochromatin, the condensed and repressive state of chromatin, represents an essential gene regulatory, organizational and architectural principle of eukaryotic genomes. Its key function is to ensure genome integrity by restricting the activity of transposable elements, preventing illegitimate recombination within repetitive genomic sequences, and supporting chromosome segregation (Fedoroff, 2012; Grewal and Moazed, 2003; Janssen et al., 2018). Given its essential roles and its strong inhibitory impact on transcription, the efficient yet specific establishment of heterochromatin is crucial.

Establishing heterochromatin requires enzymes that modify histone tails (primarily histone deacetylation and Histone 3 lysine 9 methylation) and effector proteins that recognize these specific chromatin marks and whose activity leads to chromatin compaction, decreased nucleosome turnover, and transcriptional repression (reviewed in Allshire and Madhani, 2018). To direct the general heterochromatin machinery to defined genomic loci, cells use a variety of sequence specific strategies. Besides pathways that target sequence motifs in DNA (Ninova et al., 2019; Yang et al., 2017), an alternative and highly adaptive principle relies on small regulatory RNAs that guide nuclear Argonaute proteins to complementary nascent transcripts on chromatin (Grewal, 2010; Martienssen and Moazed, 2015).

The main nuclear small RNA pathway in metazoans is the PIWI-interacting RNA (piRNA) pathway (Czech et al., 2018; Ozata et al., 2018; Siomi et al., 2011). It operates primarily in gonads and protects the germline genome from invading transposons. In *Drosophila*, the nuclear Argonaute Piwi (Cox et al., 2000) targets nascent transposon transcripts by virtue of sequence complementarity to its associated piRNAs (Brennecke et al., 2007; Saito et al., 2006; Vagin et al., 2006). By poorly understood mechanisms, this leads to heterochromatin formation and potent repression of transcription at piRNA target loci (Le Thomas et al., 2013; Rozhkov et al., 2013; Sienski et al., 2012; Wang and Elgin, 2011).

To mediate co-transcriptional silencing, Piwi requires a multitude of nuclear factors, which can broadly be divided into two categories: Group I proteins (in flies: Asterix/Gtsf1, Maelstrom, and the SFiNX complex) are piRNA pathway specific. Their molecular functions are largely unknown, but their loss specifically leads to defects in Piwi-mediated heterochromatin formation and transposon de-repression in gonads. Consequently, flies with mutations in group I factors are sterile but viable (Batki et al., 2019; Donertas et al., 2013; Eastwood et al., 2021; Fabry et al., 2019; Muerdter et al., 2013; Murano et al., 2019; Ohtani et al., 2013; Onishi et al., 2020; Schnabl et al., 2021; Sienski et al., 2015; Sienski et al., 2012; Yu et al., 2015). Group II proteins are also required for piRNA-guided heterochromatin formation, yet they execute heterochromatin formation downstream of various processes that specify heterochromatin. Group II factors are expressed in all tissues, their loss results in lethality, and they can therefore be classified as components of the general heterochromatin machinery. Examples of group II factors required for Piwi-mediated silencing are H3K9 methyltransferases, H3K4 demethylases, histone deacetylases, chromatin remodelers, the SUMO pathway, and proteins involved in chromatin compaction (Iwasaki et al., 2016; Mugat et al., 2020; Ninova et al., 2020a; Osumi et al., 2019; Sienski et al., 2015; Yu et al., 2015; Yang et al., 2019). When experimentally recruited to a reporter transgene via a heterologous DNA-binding domain, several group II factors are sufficient to initiate heterochromatin formation and transcriptional silencing (Batki et al., 2019; Ninova et al., 2020a; Yang et al., 2019). However, the mechanistic basis of how, and in which order, the piRNA pathway connects to group II factors, and how the underlying molecular interactions are controlled, is unknown.

Within the *Drosophila* nuclear piRNA-pathway, the dimeric SFiNX complex (consisting of Panoramix, the Nxf2–Nxt1 heterodimer, and the dimerization factor LC8/Cutup) is the prime candidate for a piRNA pathway-specific factor acting at the interface to the heterochromatin machinery (Batki et al., 2019; Eastwood et al., 2021; Fabry et al., 2019; Murano et al., 2019; Schnabl et al., 2021; Sienski et al., 2015; Yu et al., 2015; Zhao et al., 2019). First, SFiNX is genetically required for Piwi-piRNA complexes to silence their targets. Second, experimental tethering of SFiNX to a nascent transcript induces co-transcriptional silencing and heterochromatin formation, independent of Piwi and group I factors. SFiNX is the only known piRNA pathway factor capable of inducing robust silencing, though Maelstrom tethering can induce silencing in some reporter constellations (Onishi et al., 2020). SFiNX’s ability to induce silencing relies on the subunit Panoramix (Panx), an orphan protein with no similarity to known proteins or protein domains. Here we show that Piwi-mediated stabilization of SFiNX on chromatin leads to the multi-site conjugation of Panx with SUMO, a ubiquitin-like modifier (Gareau and Lima, 2010; Geiss-Friedlander and Melchior, 2007; Jentsch and Psakhye, 2013). SUMOylation of Panx’s disordered silencing domain enables its direct interaction with the zinc finger repressor Small ovary (Sov), which is required for piRNA-guided, as well as for global heterochromatin formation (Benner et al., 2019; Czech et al., 2013; Jankovics et al., 2018; Ninova et al., 2020b). Our work uncovers the molecular principle that connects the piRNA pathway, once engaged at a target site, to the heterochromatin machinery.

## RESULTS

### An amphipathic LxxLL motif in the intrinsically disordered silencing domain of Panx binds Sov

To understand the molecular mechanisms underlying SFiNX-mediated heterochromatin formation, we focused on Panx, the subunit that encompasses SFiNX’s silencing capacity. In cultured ovarian somatic stem cells (OSCs), Gal4-UAS mediated recruitment of Panx upstream of a GFP reporter transgene resulted in ∼25-fold repression of GFP levels (Figure 1A, B). Based on amino acid composition and predictions for protein disorder and secondary structure, we divided Panx into three parts (Figure 1C): An acidic, proline-rich and intrinsically disordered N-terminal region (IDR; aa 1-195), an NLS-containing and positively charged central region (NCR; aa 196-262), and a mostly structured C-terminal part (aa 263-541), which interacts with Nxf2–Nxt1 and Cut up (Batki et al., 2019; Eastwood et al., 2021; Fabry et al., 2019; Murano et al., 2019; Schnabl et al., 2021; Zhao et al., 2019). Of the three Panx regions, the IDR harbored strong silencing capacity (Figure 1B). We attributed the more potent repressor activity of the IDR compared to full length Panx to its higher expression levels (Figure S1A), and the residual activity of the structured C-terminus to its dimerization with endogenous, full-length Panx (Eastwood et al., 2021; Schnabl et al., 2021). As for full-length Panx, IDR-mediated silencing was accompanied by H3K9 tri-methylation and hence heterochromatin formation at the reporter locus (Figure S1B). These experiments defined the acidic IDR as the critical silencing domain within Panx.

**Figure 1:**
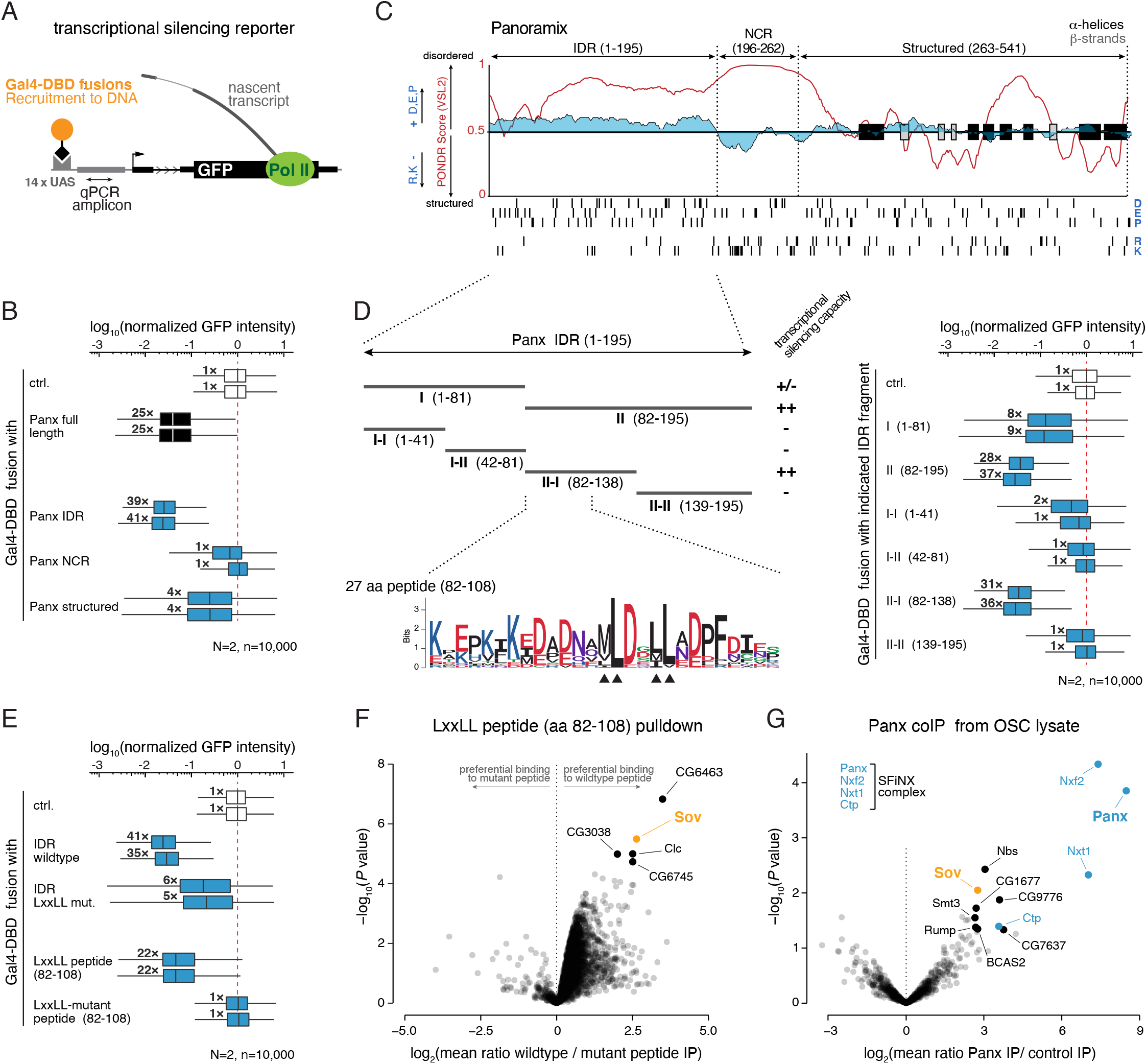
An amphipathic LxxLL motif in the Panx IDR binds Sov. **A**, Schematic representation of the GFP reporter assay in OSCs that allows for UAS - Gal4-DBD (DNA binding domain) mediated recruitment of proteins of interest upstream of the reporter transcription start site (TSS). qPCR amplicon for Figure S1B is indicated. **B**, Boxplots showing GFP reporter levels in OSCs following transfection with plasmids encoding Gal4-DBD fusions of Panx or the indicated Panx fragments (numbers indicate median fold-change, normalized to median GFP fluorescence of cells transfected with Gal4-only expressing plasmid in two biological replicates, *n* = 10,000; box plots indicate median (center line), first and third quartiles (box), whiskers show 1.5× interquartile range; outliers were omitted). **C**, Cartoon of the Panx primary sequence, indicating secondary structure elements (black, grey), protein disorder score (red; based on protein disorder predictor PONDR VSL2) and occurrence of D, E, P (positive) and K, R (negative) residues (blue line and instances indicated at bottom). IDR (intrinsic disorder region), NCR (NLS containing region) and structured region are indicated. **D**, To the left, Panx IDR fragments tested in the transcriptional silencing reporter assay are shown. The protein sequence logo shown below illustrates the pattern of amino acid conservation in the 27 amino acid peptide surrounding the conserved LxxLL motif (logo based on a multiple sequence alignment of Panx orthologs of the ‘melanogaster’ subgroup; residues colored by chemical properties-hydrophobic in black, basic in blue, acidic in red, neutral in purple, and polar in green). To the right: As in panel B, with indicated Gal4-DBD fusions. **E**, As in panel B, with indicated Gal4-DBD fusions. **F**, Volcano plot showing fold enrichment of proteins determined by quantitative mass spectrometry in Panx LxxLL-peptide pulldown experiments versus Panx-LxxLL mutant peptide control (*n* = 3 biological replicates; p-values corrected for multiple testing; Doblmann et al., 2018). **G**, Volcano plot showing fold enrichment of proteins determined by quantitative mass spectrometry in GFP-FLAG-Panx co-immunoprecipitates versus control experiments (*n* = 4 biological replicates; p-values corrected for multiple testing).

To narrow down the silencing activity within the Panx IDR, we recruited sub-fragments to the reporter locus (Figure 1D left; Figure S1C). This revealed a strong repressor activity within a ∼50 amino acid polypeptide (aa 82-138) that harbors a conserved hydrophobic motif (<u>ML</u>DS<u>LL</u>) (Figure 1D). The MLDSLL motif is reminiscent of the LxxLL motif, known from transcriptional regulators due to its role in interacting with co-activators and repressors (Plevin et al., 2005). Mutating the three leucine residues of the MLDSLL motif (MNDSQQ variant) greatly reduced, but did not abolish, the silencing capacity of the full IDR (Figure 1E; Figure S1D). However, in the context of a 27 amino acid peptide (aa 82-108), which has strong silencing capacity on its own, mutation of the LxxLL motif abrogated all repressive activity (Figure 1E; Figure S1D). Thus, the LxxLL motif is an important, but not the only, silencing determinant of Panx.

To identify factors that bind the Panx LxxLL motif, we coupled biotinylated peptides (aa 82-108) harboring the wildtype or the mutant motif to streptavidin beads and performed pulldown experiments with OSC nuclear extract. Quantitative mass spectrometry revealed a handful of significantly enriched proteins (Figure 1F; Table S1). Among the top interactors was the zinc finger protein Small ovary (Sov). Sov is essential for viability, required for transposon silencing, localizes to and is required for heterochromatin formation and interacts with Heterochromatin Protein 1 (HP1; Su(var)205) (Benner et al., 2019; Czech et al., 2013; Jankovics et al., 2018; Ninova et al., 2020b). In support of a physical Panx–Sov interaction, immuno-precipitation of GFP-tagged Panx from nuclear OSC lysate resulted in the specific co-purification of Sov (Figure 1G; Table S1). Together, our findings suggest that Panx, likely via an amphipathic LxxLL motif in its IDR, interacts with the general heterochromatin factor Sov.

### Sov is required for Piwi and Panx-mediated heterochromatin formation

To test whether the identified physical interaction between Panx and Sov is functionally relevant, we turned to a co-transcriptional silencing assay that mimics piRNA-guided repression *in vivo*. Here, aptamer-based recruitment of Panx, specifically in germline cells, via the *λ*N–boxB system to nascent transcripts of a GFP-reporter results in silencing through heterochromatin formation (Figure 2A, B) (Sienski et al., 2015; Yu et al., 2015). Depletion of Sov, via transgenic RNAi in the germline, abolished GFP-silencing, indicating that Panx requires Sov for co-transcriptional silencing (Figure 2B; Figure S2A). To extend these findings to endogenous Panx targets we turned to OSCs, where piRNA-guided silencing and heterochromatin formation at transposon loci can be most accurately studied. We first determined H3K9me3 profiles in control or Sov-depleted OSCs. H3K9me3 levels within piRNA-targeted transposons (e.g. the endogenous retroviruses *gypsy* and *mdg1*) as well as in genomic regions flanking piRNA-repressed transposon insertions (mapped in the OSC genome) were strongly reduced in cells lacking Sov (Figure 2C; Figure S2B). In line with this, transposons under Piwi control in OSCs (as defined in Sienski et al., 2012) were strongly de-repressed in Sov-depleted cells (Figure 2D). Consistent with a direct involvement of Sov at piRNA-targeted transposons, ChIP-Seq experiments using an OSC line expressing endogenously GFP-tagged Sov revealed an enrichment of Sov at piRNA-targeted transposons and the genomic regions flanking the insertions of these transposons (Figure 2E; Figure S2C). Finally, we asked whether recruiting Sov ectopically to chromatin is sufficient to establish heterochromatin. Using the DNA tethering system in OSCs (Figure 1A), we targeted Sov upstream of the GFP reporter transgene. Four days after transfecting the Gal4-Sov expressing plasmid, we observed strong reporter silencing accompanied by increased H3K9me3 levels (Figure 2F, G; Figure S2D). Our data confirm and extend previous findings that Sov is critically involved in Piwi and Panx-mediated transposon silencing (Benner et al., 2019; Jankovics et al., 2018). However, Sov does not act exclusively within the piRNA pathway: unlike piRNA pathway factor mutants, *sov* null mutants are lethal and loss of Sov impacts general heterochromatin formation (Benner et al., 2019; Jankovics et al., 2018; Ninova et al., 2020b). In support of this, depletion of Sov in OSCs mimicked depletion of the general heterochromatin factor HP1 and led not only to de-silencing of Piwi-repressed transposons, but also of numerous other transposons not impacted by loss of Piwi or Panx (e.g. *G6* or *gypsy7*; Figure 2D, H). Based on these findings, we concluded that the physical connection between Panx and Sov is a major intersection point between piRNA pathway and general heterochromatin machinery. We therefore set out to molecularly dissect the Panx–Sov interaction.

**Figure 2:**
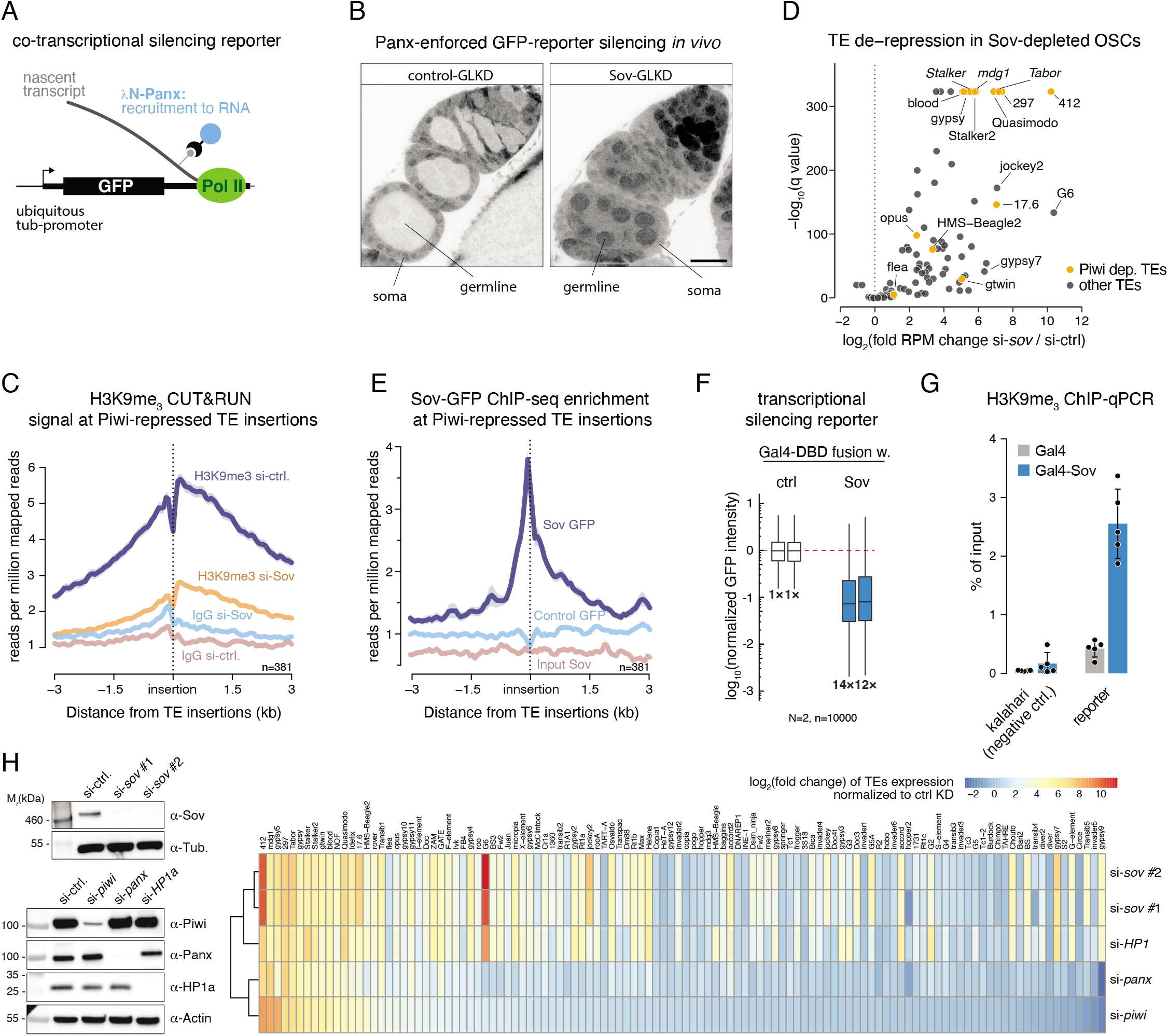
Sov is required for Piwi and Panx-mediated heterochromatin formation. **A**, Schematic representation of the GFP silencing reporter in flies, which allows for recruitment of *λ*N-tagged proteins to the nascent transcript via boxB sites in the 3’ UTR. **B**, Confocal images of early oogenesis stages showing fluorescence levels (greyscale; scale bar: 20μm) of the ubiquitously expressed GFP reporter with *λ*N-Panx expressed in all germline cells and additional germline-specific knockdown (GLKD) against *white* (control; left) or *sov* (right). **C**, Metaplot of H3K9me3 levels (in OSCs) at regions flanking piRNA-targeted transposon insertions (vertical line) following depletion of Sov as measured by Cut&Run (*n* = 381 transposon insertions). **D**, Volcano plot showing fold changes in steady state RNA levels of annotated transposons in Sov-depleted OSCs compared to control (piRNA-repressed transposons marked in yellow; *n* = 3). **E**, Metaplot of Sov-GFP enrichment (in OSCs) at regions flanking piRNA-targeted transposon insertions (vertical line) determined by ChIP-seq (*n* = 381 transposon insertions). **F**, Boxplots showing GFP reporter (Fig. 1A) levels in OSCs following transfection with plasmids encoding a Gal4-DBD fusion of Sov (numbers indicate median fold-change, normalized to median GFP fluorescence of cells transfected with Gal4-only expressing plasmid in two biological replicates, *n* = 10,000; box plots indicate median (center line), first and third quartiles (box), whiskers show 1.5× interquartile range; outliers were omitted). **G**, H3K9me3 levels at the GFP reporter locus (amplicon indicated in Fig. 1A) after Sov tethering determined by ChIP-qPCR (*n* = 5 biological replicates; the gene desert ‘*kalahari’* served as negative control; error bars: St. dev. (standard deviation)). **H**, Heatmap showing the fold change of steady state RNA levels (determined by RNA-seq) of annotated *Drosophila* transposons in OSCs after siRNA-mediated Sov, HP1, Panx or Piwi depletion (depletion shown by western blot experiments to the left).

### Structural basis of the Panx–Sov interaction

The 370 kDa Sov protein lacks annotated domains in its ∼1,500 amino acid N-terminal half and harbors 21 C2H2 zinc finger domains in its C-terminal half (Figure 3A) (Benner et al., 2019; Jankovics et al., 2018). To identify the region within Sov responsible for binding to Panx, we performed a Panx LxxLL peptide pulldown using nuclear OSC lysate subjected to mild sonication, which resulted in fragmentation of the Sov protein. We then determined where the identified peptides map along the Sov primary sequence and observed a strong clustering at the Sov N-terminus (Figure 3A). As LxxLL motifs in disordered regions of transcriptional regulators often bind to α-helical domains of interacting proteins (Plevin et al., 2005), we searched for predicted folded domains in the N-terminal region of Sov with the protein homology algorithm HHPRED (Zimmermann et al., 2018). This revealed a putative α-helical domain within the first one hundred amino acids of Sov (termed N-terminal domain, NTD; Figure 3A). To test whether this domain interacts with the Panx LxxLL peptide, we co-expressed GFP-tagged Sov NTD (aa 1-118) and the Panx LxxLL peptide (aa 82-108) in Schneider cells, which lack a piRNA pathway. Co-immunoprecipitation experiments revealed that the wildtype Panx peptide (fused to Gal4_DBD-FLAG), but not the peptide with mutated LxxLL motif, interacted with the Sov NTD but not with GFP alone (Figure 3B). Similarly, when coupled to streptavidin beads, a biotinylated Panx peptide with LxxLL motif, but not the mutant variant, interacted with recombinant Sov NTD, whose extent could be refined to residues 14-90 (Figure 3C). Based on isothermal calorimetry (ITC) measurements, the Sov NTD bound the Panx LxxLL peptide with a dissociation constant of 0.87 ± 0.39 μM, while the mutant peptide did not show measurable binding (Figure 3D). These findings are in line with our previous observation that the Panx peptide (aa 82-108) harboring the mutated LxxLL motif was inert in the reporter silencing assay (Fig. 1E).

**Figure 3:**
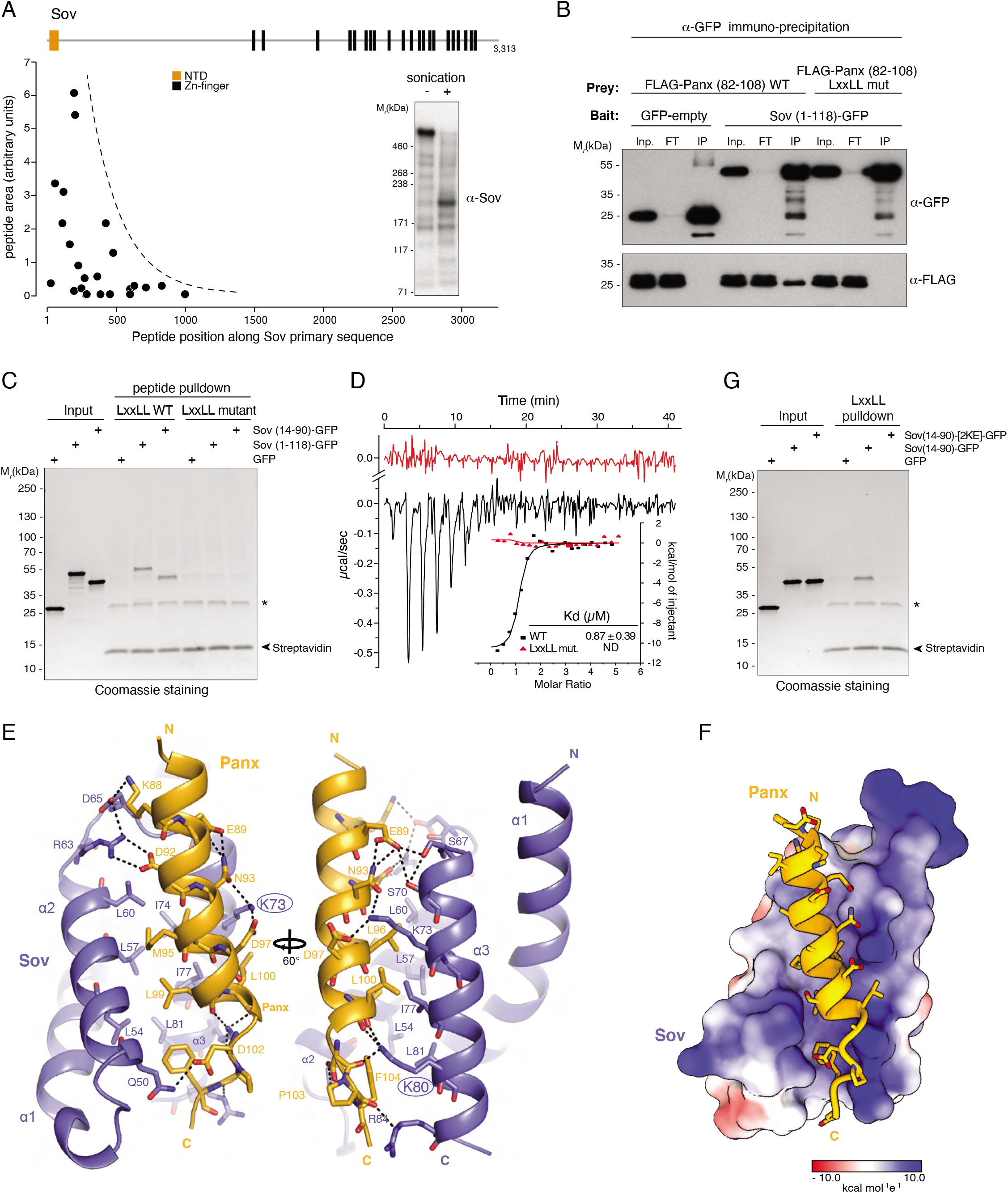
Structural basis of the Panx–Sov interaction. **A**, Shown is the distribution (along the Sov primary sequence; annotated domain organization at the top) and relative level of Sov peptides identified by mass spectrometry from a Panx LxxLL peptide pulldown. The western blot inlay to the right indicates Sov protein integrity upon sonication of nuclear OSC lysate. **B**, Western blot analysis of immunoprecipitation experiments from S2 cells transiently expressing GFP-Sov (aa 1-118) as bait and wildtype or mutant Gal4-DBD_FLAG-tagged Panx LxxLL peptide (aa 82-108) as prey. **C,** Coomassie stained SDS-PAGE showing an *in vitro* pulldown experiment with streptavidin-bound wildtype or mutant Panx LxxLL peptide and recombinant GFP-tagged Sov NTD fragments as prey (asterisk indicates a background band from Streptavidin beads). **D**, Isothermal calorimetry measurement of the interaction affinity of the Sov NTD (aa 14-90) with wildtype (black) or mutant (red) Panx LxxLL peptide (aa 82-108). **E**, Shown are ribbon models of the Sov NTD (aa 14-90; blue) - Panx LxxLL peptide (aa 82-108; gold) structure with interacting residues in bonds representation (K73 and K80 residues mutated in panel G are highlighted). **F,** Surface representation of the Sov NTD colored according to electrostatic surface potential (red, negative; white, neutral; blue, positive) bound to the Panx LxxLL peptide (gold) as ribbon model with sidechains shown in bonds representation.

To gain atomic insight into the Panx–Sov interaction, we determined the X-ray crystal structure of the Sov NTD (aa 14-90) bound to the Panx peptide (aa 82-108) at 2.5 Å resolution. The Sov NTD folds into a three-helix bundle with helices α2 and α3 directly contacting the Panx LxxLL peptide, which adopts an α-helical conformation (Figure 3E). Two types of interactions underlie the specific Panx-Sov association: first, a hydrophobic cleft within the Sov NTD formed by α2 (residues L54, L57, L60) and α3 (residues I74, I77, L81) accommodates the hydrophobic LxxLL motif (residues M95, L96, L99, L100) of the Panx helix (Figure 3E). Second, the acidic Panx helix engages in hydrogen bond and salt bridge interactions with several positively charged residues lining the hydrophobic cleft of the Sov NTD (Figure 3E, F). To experimentally test the structural findings, we purified a mutant Sov NTD predicted to be incompatible with Panx binding. A charge reversal of two solvent accessible residues that do not contribute to stabilize the overall Sov NTD fold (K73E, K80E), resulted in loss of interaction with the Panx LxxLL peptide (Figure 3E, G). Altogether, our biochemical and structural data demonstrate a direct protein-protein interaction between the piRNA pathway factor Panx and the general heterochromatin factor Sov.

### A dual-binding mode between Panx and Sov, coordinated by Panx SUMOylation

Since Panx requires Sov for silencing (Figure 2), we hypothesized that mutating the LxxLL motif within Panx should result in female sterility as seen in *panx* mutant flies. However, flies expressing a Panx variant with the M<u>N</u>DS<u>QQ</u> mutation showed only moderate fertility defects (∼70% embryo hatching rate). Deleting the entire IDR of Panx instead resulted in complete sterility, phenocopying *panx* null mutants (Figure 4A). This mirrored results from the heterologous tethering assay, in which the LxxLL mutant IDR retained moderate silencing activity (Figure 1E) and suggested additional silencing determinants or LxxLL-independent physical links between Panx and Sov. We focused on Smt3 (*Drosophila* SUMO), a small Ubiquitin-like protein that has been linked to Piwi-mediated silencing (Ninova et al., 2020a), and that was enriched in co-immunoprecipitation experiments from OSC lysate using both, Panx or Sov as bait (Figure 1G; Figure S3A; Table S1). SUMO is covalently conjugated to acceptor lysine residues (often within the consensus sequence Ψ-K-x-D/E, with Ψ being a large hydrophobic residue) in accessible regions of substrate proteins and can mediate or strengthen protein-protein interactions (Gareau and Lima, 2010; Jentsch and Psakhye, 2013; Pichler et al., 2017). This requires one binding partner to be SUMOylated and the other to contain a SUMO interacting motif (SIM). When analyzing the primary sequences of Panx and Sov, we found an intriguing pattern that would be consistent with a SUMO-dependent Panx–Sov interaction: Five of the eleven lysine residues within the Panx IDR are predicted SUMOylation sites (K6, K10, K82, K88, K143). The Sov NTD on the other hand is flanked on both sides by two SIMs (computational predictions after Zhao et al., 2014).

**Figure 4:**
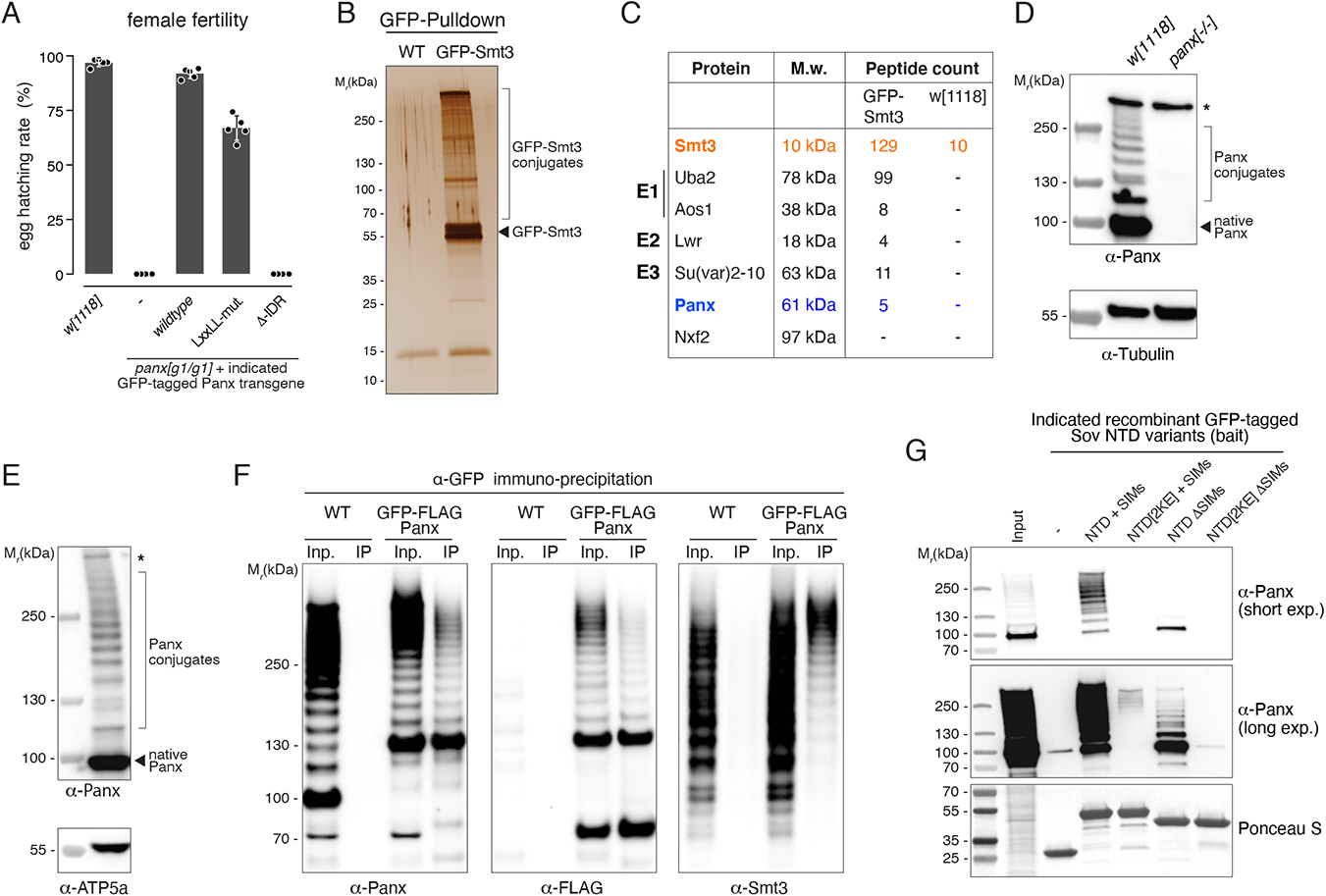
Panx is SUMOylated. **A**, Hatching rates of eggs laid by females with indicated genotype mated to wildtype males (*n* = 5; error bars: St. dev.). **B**, Silver stained SDS-PAGE of a pulldown experiment (with denaturing wash steps) using GFP nanobodies and ovarian lysate from GFP-Smt3 expressing flies or control flies. **C**, Unique peptide counts of indicated proteins identified by mass spectrometry in the pulldown experiment shown in panel B. **D, E,** Western blot analysis of ovary lysates (panel D) or OSC lysate (panel E) probed with an α-Panx antibody (asterisk indicates an unspecific band; native Panx runs at ∼100 kDa despite a molecular weight of 61 kDa). **F,** Western blots showing GFP-Trap immunoprecipitation experiments with lysates from wildtype (WT) OSCs or OSCs expressing endogenously GFP-FLAG-tagged Panx (Inp: input; IP: immuno-precipitate). The band at ∼70 kDa in the α-FLAG western represents an N-terminal Panx degradation product. **G,** Western blots showing a pulldown experiment using indicated recombinant GFP-tagged Sov NTD variants as bait and nuclear OSC lysate as input.

To investigate whether Panx is SUMOylated *in vivo*, we generated flies expressing GFP-tagged Smt3 from the *smt3* promoter, and immuno-precipitated GFP-Smt3 from ovarian lysate under denaturing wash conditions, thereby enriching for proteins covalently bound to Smt3 (Figure 4B; Figure S3B; Table S1). Among the enriched proteins (compared to an IP from wildtype ovary lysate) were the core SUMOylation machinery (Smt3, E1 activating enzyme Uba2–Aos1, E2 conjugating enzyme Lwr/Ubc9), the SUMO E3 Ligase Su(var)2-10, and Panx (Figure 4C). No other SFiNX subunit, nor Piwi, were enriched in the GFP-Smt3 IP. To substantiate these findings, we performed western blot experiments on whole cell extracts from ovaries and OSCs that were prepared in the presence of N-ethylmaleimide (NEM), an irreversible inhibitor of de-SUMOylating enzymes. This revealed, besides native Panx (running at ∼100 kDa despite a molecular weight of 61 kDa), a prominent ladder of Panx isoforms with increasing molecular weight, consistent with an increasing number of SUMO moieties (Figure 4D, E).

To probe whether the observed isoform ladder is indeed due to SUMOylation of Panx, we performed immuno-precipitation experiments using GFP-trap beads with lysates from either wildtype OSCs or OSCs expressing endogenously FLAG-GFP tagged Panx. Both input samples showed native Panx and the isoform ladder (Figure 4F left), and we observed an identical pattern of bands, specific for the tagged Panx cell line, with an antibody against the FLAG epitope (Figure 4F middle). After immunoprecipitation with anti-GFP nanobodies, native Panx and the higher molecular weight isoforms were specifically recovered from OSCs expressing FLAG-GFP-Panx. Western blot analysis with an antibody against Smt3 confirmed that the laddered signal represented SUMOylated Panx (Figure 4F right). Considering the absence of SUMO chains in *Drosophila* (Urena et al., 2016), our results suggest that, at steady state, a substantial fraction of Panx in OSCs is conjugated with SUMO on multiple lysine residues.

SUMO promotes protein-protein interactions by binding to SIMs in partner proteins. The best characterized SIMs are composed of three to four exposed aliphatic amino acids (I, V, L), often flanked by negatively charged residues, that bind to a hydrophobic groove in SUMO (Kerscher, 2007). The Sov core NTD (aa 14-90) is immediately flanked by two putative SIMs: EDD<u>VVVV</u> (aa 5-11) and <u>II</u>DI (aa 96-99). To test whether these predicted SIMs are relevant for the Panx–Sov interaction, we immobilized a series of recombinant Sov NTD variants on beads and incubated them with nuclear OSC lysate (Figure 4G). The core NTD without SIMs (NTD Δ-SIMs) was indifferent in terms of binding native or SUMOylated Panx. In contrast, the NTD with both flanking SIMs (NTD + SIMs) had a strong binding preference for SUMOylated Panx isoforms. An NTD that was unable to interact with the Panx LxxLL peptide but that was flanked by both SIMs (NTD[2KE] + SIMs) bound exclusively to heavily SUMOylated Panx isoforms, albeit weakly. The 2KE mutant NTD without flanking SIMs (NTD[2KE] Δ-SIMs) was inert in respect to Panx binding, comparable to GFP alone (Figure 4G). These results strongly suggest a model where SUMOylated Panx interacts with Sov via a dual mode: (1), the NTD–LxxLL interaction and (2) immediately flanking SUMO-SIM interactions.

To test the dual-binding model, we first asked whether expression of the isolated Sov N-terminus interferes in a dominant negative manner with the ability of Panx to recruit endogenous Sov. Indeed, expression of the NTD with flanking SIMs, but not of the core NTD alone, resulted in de-repression of piRNA-targeted transposons (e.g. *mdg1* and *gypsy*) and of the *expanded* gene, which is repressed in a Piwi-dependent manner due to an intronic *gypsy* insertion (Figure 5A; Figure S4A) (Sienski et al., 2012). As a more direct test, we used the DNA tethering assay (Figure 1A) and determined the repressive activity of Panx IDR variants mutated for either the LxxLL motif or the five consensus SUMOylation sites (charge-preserving lysine to arginine mutations; Panx[5KR]), or both together (Figure 5B). Both single mutant IDR constructs showed substantially reduced silencing ability but were not inert. The double mutant, however, lost all silencing capacity (Figure 5B; Figure S4B).

**Figure 5:**
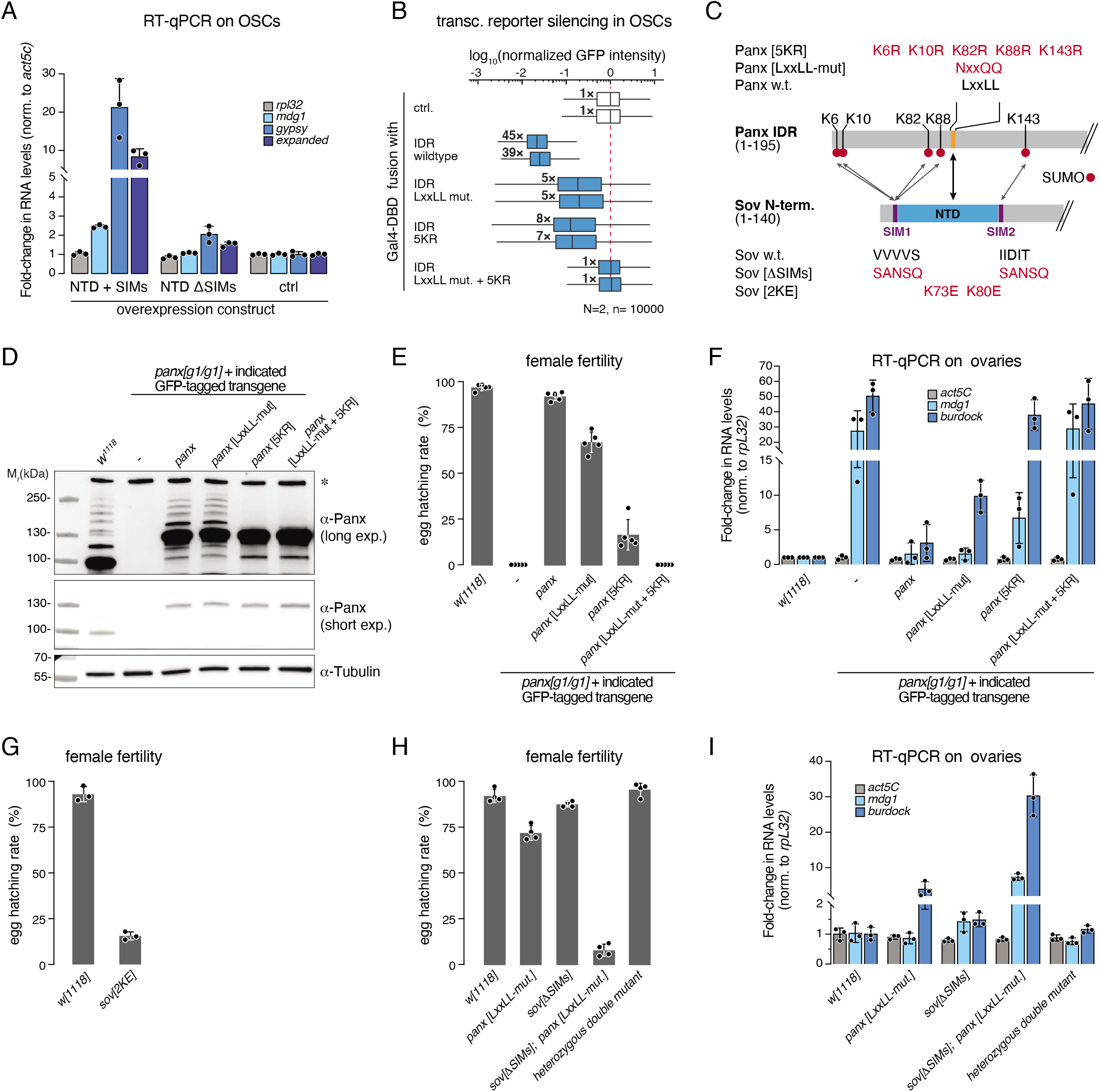
A SUMOylation-dependent dual mode interaction between Panx and Sov. **A**, RT-qPCR analysis showing fold changes in steady state RNA levels of indicated transposons in OSCs transiently overexpressing GFP-tagged Sov NTD including or excluding the flanking SIMs (*n* = 3 biological replicates; error bars: St. dev.). **B**, Boxplots of GFP intensity in OSCs expressing the transcriptional silencing reporter (Fig. 1A) following transfection with plasmids encoding Gal4-DBD fusions of the indicated Panx IDR variants (numbers indicate median fold-change, normalized to median GFP fluorescence of cells transfected with Gal4-only expressing plasmid in two biological replicates, *n* = 10,000; box plots indicate median (center line), first and third quartiles (box), whiskers show 1.5× interquartile range; outliers were omitted). **C,** Schematic representation of the SUMOylation-dependent dual interaction between Panx IDR and Sov N-terminus (identity of used Panx and Sov mutants is indicated). **D**, Western blot showing levels and SUMOylation extent of endogenous Panx or GFP-tagged Panx rescue variants expressed in fly ovaries of indicated genotype (asterisk: unspecific band). **E**, Hatching rates of eggs laid by females with indicated *panx* genotype mated to wildtype males (*n* = 5 biological replicates; data from a common experiment with Fig. 4A; error bars: St. dev.). **F**, RT-qPCR analysis showing fold changes in steady state RNA levels of indicated transposons in ovaries from flies of indicated genotype (*n* = 3 biological replicates; error bars: St. dev.). **G, H**, Hatching rates of eggs laid by females with indicated genotype mated to wildtype males (*n* = 3 or 4 biological replicates; error bars: St. dev.) **I**, RT-qPCR analysis showing fold changes in steady state RNA levels of indicated transposons in ovaries from flies of indicated genotype (*n* = 3 biological replicates; error bars: St. dev.).

Based on our biochemical and OSC experiments, we set out to genetically uncouple Panx and Sov in flies (Figure 5C). We generated transgenic fly lines expressing, instead of the endogenous protein, FLAG-GFP-tagged wildtype Panx, the single mutants Panx[LxxLL-mut] or Panx[5KR], and the double mutant Panx[LxxLL-mut+5KR]. Western blot analysis of ovarian lysates showed that the tagged proteins were expressed at levels comparable to endogenous Panx, that N-terminal tagging did not interfere with SUMOylation, and that the 5KR mutation, but not the LxxLL mutation, prevented detectable Panx SUMOylation (Figure 5D). Individually, the LxxLL mutation and the 5KR mutation had intermediate (∼67% fertile) and substantial (∼16% fertile) impact on fertility and transposon repression, respectively (Figure 5E, F). The double mutant, however, was entirely sterile and phenocopied *panx* null mutants in terms of transposon de-silencing and the collapsed morphology of laid eggs (Figure 5E, F).

We finally tested the Panx–Sov dual-binding model from the side of Sov, which unlike Panx or the piRNA pathway is required for viability. We hypothesized that the SIM-NTD-SIM unit at the Sov N-terminus represents a binding module specific for the piRNA pathway. If true, targeted mutations of this unit should uncouple Sov specifically from the piRNA pathway, yielding viable yet sterile flies. Using CRISPR-mediated genome engineering, we generated two defined *sov* alleles (Figure 5C). First, the NTD[2KE] mutant that is unable to bind the Panx LxxLL peptide *in vitro* (Figure 3G) and second, a Sov variant with point mutations in both SIMs. Females homozygous for the *sov[NTD_2KE]* allele were viable and laid eggs yet displayed strongly reduced fertility (egg hatching rate ∼15%; Figure 5G). Their sterility was more severe compared to females expressing Panx[LxxLL-mut] (egg hatching rate ∼70%), suggesting that the Sov NTD binds, besides Panx, to additional client proteins. Flies homozygous for the *sov[ΔSIM]* allele were barely affected in their fertility and ability to silence transposons (Figure 5H, I). This prompted us to generate Panx–Sov binding deficient flies where the SUMO–SIM interactions were prevented via the *sov[ΔSIM]* allele, and the LxxLL–NTD interaction via the *panx[LxxLL-mut]* allele. While both alleles individually showed a fertility of ∼72% and ∼85%, respectively, their combination resulted in almost complete sterility (hatching rate ∼8%), accompanied by strong transposon de-repression (Figure 5H, I). Our combined data show that Panx engages the general heterochromatin factor Sov via two distinct molecular interactions, and that their combined action confers strong transcriptional silencing activity to the Panx IDR, and therefore SFiNX, *in vivo*.

### SUMOylation of Panx is coupled to its Piwi-mediated stabilization on chromatin

The 5KR mutation in the IDR prevents Panx SUMOylation *in vivo* and is a strong loss of function allele despite harboring the Sov-interacting LxxLL motif (Figure 5). We hypothesized that Panx SUMOylation serves a regulatory function to gate a functional interaction between Panx and Sov. If SUMOylation of Panx would occur only at piRNA target sites, this would allow cells to control the link between piRNA pathway and general heterochromatin machinery. To test this, we reasoned that piRNA-guided silencing occurs co-transcriptionally and that nascent target transcripts are attached to chromatin via transcribing RNA Polymerase II. We therefore separated a whole cell OSCs lysate, in the presence of NEM, into a soluble fraction (enriched for tubulin) and an insoluble pellet fraction (enriched for histone H3 and therefore chromatin) (Figure 6A). Western blot analysis showed that the SUMOylated Panx isoforms were almost exclusively present in the chromatin-enriched pellet fraction, while soluble Panx was non-modified (Figure 6A). This supported a model where cells restrict SUMOylation of Panx to chromatin.

**Figure 6:**
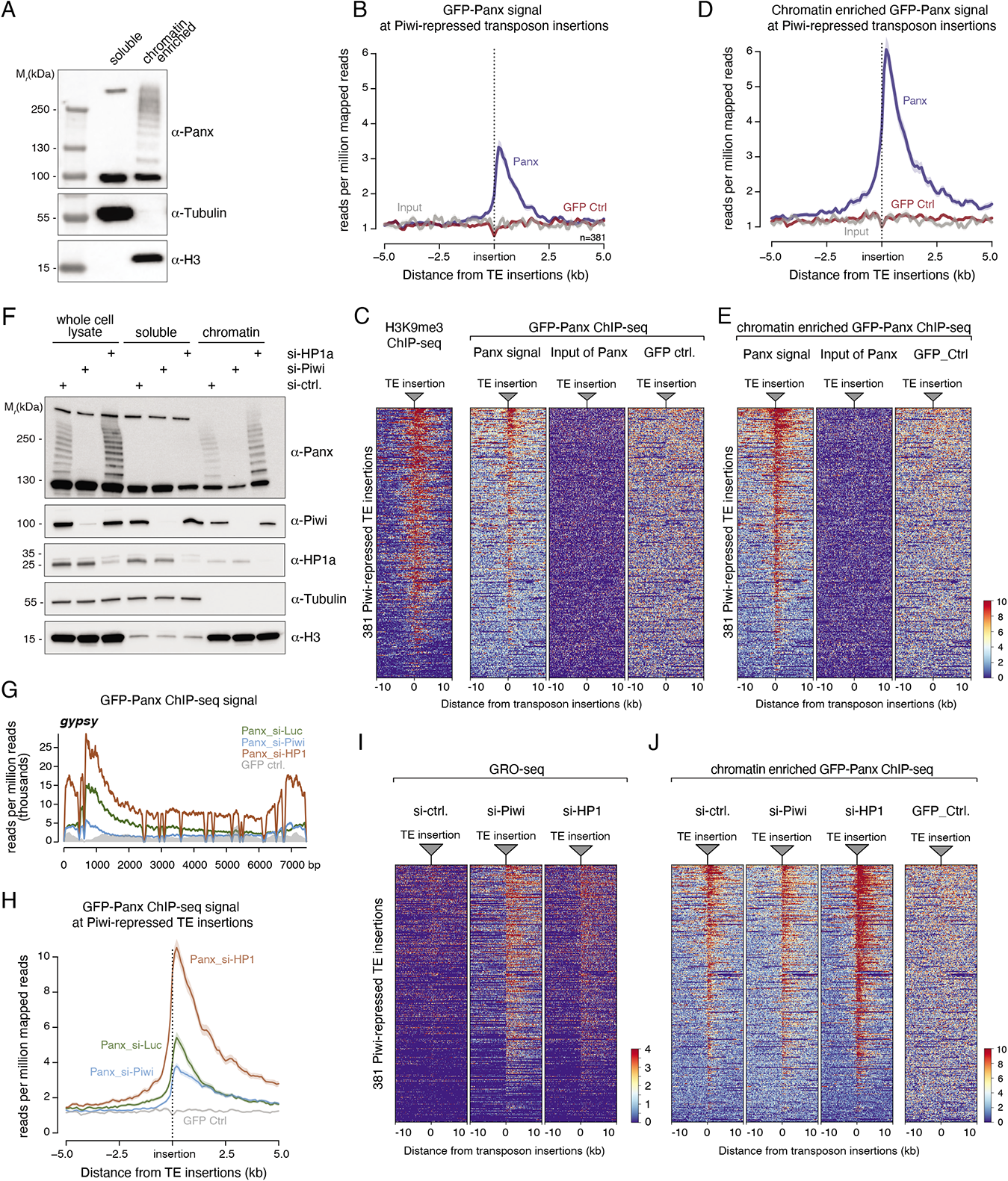
SUMOylation of Panx is coupled to its Piwi-mediated stabilization on chromatin. **A**, Western blot analysis of soluble and insoluble (chromatin-enriched) fractions from OSCs. **B**, Meta profiles of GFP-Panx enrichment at genomic regions flanking piRNA-targeted transposon insertions (vertical line) in OSCs, determined by anti-GFP ChIP-seq using OSCs expressing endogenously GFP-FLAG-tagged Panx (*n* = 381 transposon insertions). **C**, Heatmap corresponding to the meta profile in panel B. Transposon insertions were ranked by H3K9me3 signal intensity in genomic regions flanking the insertions (left). **D**, As in panel B, but ChIP experiment used pre-extracted OSCs as input. **E**, Heatmap corresponding to meta profile in panel D. **F**, Western blot analysis of whole cell, soluble and insoluble (chromatin-enriched) fractions from OSCs depleted for indicated factors via siRNA transfections. **G**, Occupancy of Panx on the *gypsy* transposon, determined via ChIP-seq using pre-extracted OSCs depleted for indicated factors. **H**, Meta profile of GFP-Panx enrichment at genomic regions flanking piRNA-targeted transposon insertions (vertical line) in OSCs depleted for indicated factors, determined by anti-GFP ChIP-seq using pre-extracted OSCs expressing endogenously GFP-FLAG-tagged Panx (*n* = 381 transposon insertions; GFP Ctrl. from a common experiment with panel D). **I**, Heatmap showing GRO-seq signal at genomic regions flanking 381 piRNA-targeted transposon insertions (vertical line) in OSCs depleted for indicated factors. **J**, Heatmap corresponding to meta profile in panel H (transposon insertion coordinates ranked by GRO-Seq signal after Piwi depletion).

To characterize the genomic binding sites of Panx, we performed ChIP-seq experiments using anti-GFP antibodies and OSCs expressing endogenously GFP-tagged Panx. Non-tagged cells served as control. This revealed an enrichment of Panx at piRNA-targeted transposons (e.g. *gypsy*) but not at non-targeted transposons (e.g. *Doc*; Figure S5A). Similarly, Panx was enriched at genomic regions flanking the 381 piRNA-targeted transposon insertions in OSCs, specifically in the tagged cell line (Figure 6B, C). To our surprise, Panx was also enriched above background at expressed gene loci (which do not give rise to piRNA complementary transcripts; Figure S5B). Considering that a substantial fraction of Panx in the chromatin-enriched fraction was not SUMOylated (Figure 6A), we reasoned that native Panx might be transiently associated with transcribed loci, possibly through intrinsic sampling of nascent transcripts, and that Panx becomes stabilized and SUMOylated only at piRNA-targeted loci. To test this, we performed ChIP-seq experiments with OSCs that were pre-extracted to enrich for more stably chromatin associated, SUMOylated Panx prior to formaldehyde crosslinking. This resulted in relatively increased Panx signal at piRNA targeted transposons and genomic regions flanking piRNA-targeted transposon insertions (Figure 6D, E; Figure S5C), while the signal at transcribed gene loci decreased (Figure S5B right).

To establish a more direct link between Piwi-mediated heterochromatin formation, Panx stabilization on chromatin, and Panx SUMOylation, we experimentally decreased or increased the levels of Panx engaged in co-transcriptional silencing. We depleted OSCs of Piwi, which should block stable chromatin association of Panx at transposon loci, or of HP1, which is required for Piwi-mediated silencing (Iwasaki et al., 2016; Yu et al., 2015) and whose absence should lead to increased levels of Piwi at derepressed transposon loci. While piRNA-targeted transposons were de-repressed in both, Piwi as well as HP1 depleted cells (Figure 2H, 6I), we observed opposing impacts on Panx SUMOylation (Figure 6F): In Piwi-depleted cells, Panx SUMOylation was absent and Panx levels in the chromatin enriched fraction were reduced. In HP1-depleted cells, on the other hand, the fraction of SUMOylated Panx increased, and all SUMOylated Panx isoforms were again present in the chromatin-enriched fraction (Figure 6F). Consistent with these findings, Panx occupancy on nearly all piRNA-repressed transposons (e.g. *gypsy*) and in the genomic regions flanking piRNA-repressed transposon insertions was reduced in OSCs depleted for Piwi but was increased in OSCs depleted for HP1 (Figure 6G, H, J; Figure S5D-F). Transposons not targeted by Piwi (e.g. *Doc* or *F-*element) did not show Panx enrichment above background in either condition (Figure S5F). Taken together, our data support a model where binding of Piwi-piRNA complexes to complementary nascent transcripts results in SUMOylation and stabilization of Panx on chromatin, thereby setting the stage for recruiting the heterochromatin machinery specifically to piRNA target loci.

### Direct Ubc-9 mediated SUMOylation of Panx is independent of Su(var)2-10

The specific SUMOylation of Panx at piRNA target sites raises the question of how this process is molecularly controlled. Protein SUMOylation is an ATP dependent reaction that requires the consecutive action of E1 activating enzyme (Aos1–Uba2 in *Drosophila*) and E2 conjugating enzyme (Lwr/Ubc9 in *Drosophila*). In most cases, specific E3 ligases stimulate the E2-catalyzed conjugation of SUMO to a target lysine residue of the substrate (Gareau and Lima, 2010; Geiss-Friedlander and Melchior, 2007; Jentsch and Psakhye, 2013). We first focused on the E3 Ligase Su(var)2-10, which is genetically required for Piwi- and Panx-mediated transcriptional silencing and heterochromatin formation (Hari et al., 2001; Ninova et al., 2020a). The PIAS family protein Su(var)2-10 has been proposed to promote protein-group SUMOylation at genomic piRNA target loci (Ninova et al., 2020a), thereby creating binding platforms for heterochromatin effectors (e.g. histone methyltransferases or histone deacetylases) via multiple SUMO–SIM interactions (Jentsch and Psakhye, 2013). The substrates of Su(var)2-10 in the piRNA pathway are unknown and we therefore asked whether SUMOylation of Panx depends on Su(var)2-10. We first examined the dynamics of Panx SUMOylation upon loss of the core SUMO-pathway and depleted OSCs for Smt3 (Figure 7A). 48 or 72h after siRNA transfection (longer siRNA treatments resulted in cell death), free Smt3 was barely detectable and overall Smt3-conjugates were reduced. In line with this, SUMOylated Panx isoforms were reduced (Figure 7A). Unexpectedly however, native Panx levels were also reduced. This was markedly different to Piwi-depleted cells, where Panx SUMOylation was lost yet native Panx levels were unchanged (Figure 6F), suggesting that the inability to SUMOylate Panx stabilized at piRNA-target sites leads to its degradation. Upon siRNA-mediated depletion of the E2 conjugating enzyme Lwr/Ubc9 (Figure S6A), we similarly observed reduced levels of native Panx and a mild loss of SUMOylated Panx isoforms, especially for the 72h sample (Figure 7B). In stark contrast, depletion of Su(var)2-10 to nearly undetectable levels resulted in increased Panx SUMOylation, both 48 and 72h after siRNA transfection, and levels of native Panx were not reduced (Figure 7C). The observed increase in Panx SUMOylation might result from de-repression of piRNA-targeted transposons (leading to more Piwi-targeting at chromatin similar to the HP1 depletion) or from an increased availability of Ubc9-Smt3 conjugates in Su(var)2-10 depleted cells. We concluded that Su(var)2-10, despite its essential role in Piwi-mediated heterochromatin formation (Ninova et al., 2020a), is not required for Panx SUMOylation and therefore most likely acts more downstream.

**Figure 7:**
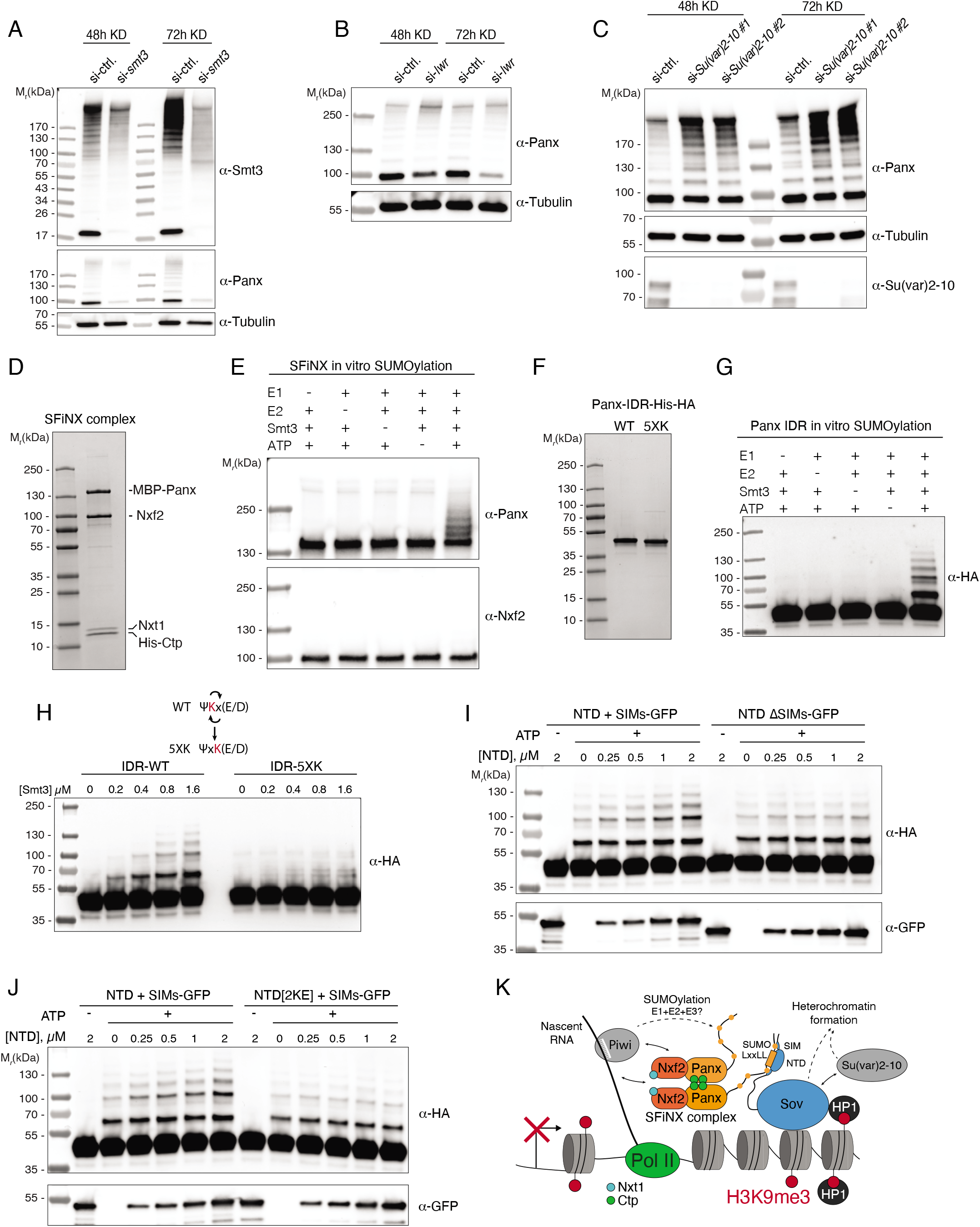
Direct Panx SUMOylation by Ubc9, independent of Su(var)2-10 and enhanced by Sov. **A**, Western blot analysis showing depletion of Smt3 and the associated decrease in SUMOylated proteins as well as SUMOylated and native Panx in OSCs. **B**, Western blot analysis showing changes in SUMOylated and native Panx in OSCs depleted for Lwr/Ubc9. **C**, Western blot analysis showing the depletion of Su(var)2-10 and the associated changes in the level of SUMOylated Panx in OSCs. **D**, Coomassie-stained SDS-PAGE showing recombinant full length SFiNX complex composed of TwinStrep-MBP-Panx, His6-Ctp, Nxf2 and Nxt1). **E**, Western blot analysis of *in vitro* SUMOylation assay with full length SFiNX complex as substrate. **F**, Coomassie-stained SDS-PAGE of recombinant WT and 5XK mutant Panx IDR-3xHA-His10. **G**, Western blot analysis of *in vitro* SUMOylation assay with WT Panx IDR as a substrate. **H**, Western blot analysis showing *in vitro* SUMOylation efficiencies (increasing concentration of Smt3) of WT and 5XK mutant Panx IDR. **I**, Western blot analysis showing the enhancement of Panx IDR *in vitro* SUMOylation by the Sov NTD in a SIM-dependent manner. **J**, Western blot analysis showing the enhancement of Panx IDR *in vitro* SUMOylation by the Sov NTD in a LxxLL binding-dependent manner. **K**, Schematic model summarizing the identity and regulation of the molecular interface between piRNA pathway (SFiNX) and general heterochromatin machinery (Sov).

Besides Su(var)2-10, only few other E3 SUMO ligases are known, and none of these were identified in genetic screens for transposon silencing factors. We therefore explored an alternative model: Unlike protein ubiquitination, where E3 ligases are required for E2-mediated substrate conjugation, SUMOylation can occur in an E3 independent manner (Gareau and Lima, 2010; Johnson and Blobel, 1997; Rodriguez et al., 2001; Sampson et al., 2001). This requires an accessible SUMOylation consensus sequence (Ψ-K-x-D/E) that directly interacts with the substrate binding groove of the E2 conjugating enzyme Ubc9 (Bernier-Villamor et al., 2002). The Panx IDR contains four Ψ-K-x-D/E consensus sites. To test whether Ubc9 can SUMOylate Panx independent of an E3-ligase, we turned to an *in vitro* SUMOylation assay (Flotho et al., 2012). We purified recombinant *Drosophila* Smt3, the dimeric E1 activating enzyme Aos1–Uba2, the E2 conjugating enzyme Lwr/Ubc9 and the complete SFiNX complex consisting of full length Panx (Strep-MBP-tagged), the Nxf2–Nxt1 heterodimer, and the Dynein light chain protein Ctp (Figure 7D; Figure S6B). When incubating all factors together for 1h at 30°C, we observed SUMOylation of Panx with up to five Smt3 moieties in an ATP, Smt3, and E1/E2 dependent manner (Figure 7E). In contrast, the SFiNX subunit Nxf2 was not modified, consistent with it lacking predicted SUMOylation sites and with the absence of higher molecular weight Nxf2 isoforms in OSCs (Figure 7E; Figure S6C).

To substantiate our findings, we tested whether E1/E2-dependent SUMOylation of Panx requires the targeted lysine residues to reside in the Ψ-K-x-D/E context. We focused on the IDR, which harbors all predicted strong SUMOylation consensus sites in Panx. We purified recombinant Panx IDR (aa 1-195) with a His-HA tag (which lacks lysine residues; Figure 7F). Like full length Panx, the Panx IDR was readily SUMOylated in an ATP, Smt3, and E1/E2 dependent manner (Figure 7G). In contrast, an IDR variant where the two central residues within each SUMOylation consensus site were swapped, thereby resulting in Ψ-x-K-D/E motifs, was a very poor SUMOylation substrate (Figure 7H). These findings would support a model where Ubc9 SUMOylates the Panx IDR in an E3-independent manner.

Protein SUMOylation can be stimulated by substrate specific E3 ligases, but also through SIMs *in cis* that coordinate Ubc9-SUMO conjugates proximal to target lysines (Lin et al., 2006; Meulmeester et al., 2008). Considering that the Sov NTD interacts with the LxxLL motif in the Panx IDR and possesses two immediately adjacent SIMs, we asked whether the Sov N-terminus might stimulate Panx SUMOylation *in trans*. Indeed, the Sov NTD flanked by two SIMs stimulated SUMOylation of the Panx IDR *in vitro* (Figure 7I). Sov NTD variants with only one flanking SIM showed weak stimulatory activity (Figure S6D). The Sov core NTD without flanking SIMs was unable to stimulate SUMOylation of the Panx IDR, pointing to a direct involvement of the two SIMs rather than a model where NTD binding opens up the Panx IDR for optimal SUMOylation (Figure 7I). Similarly, an NTD variant with two flanking SIMs yet unable to bind the Panx LxxLL motif (2KE mutant), did not stimulate SUMOylation of the Panx IDR (Figure 7J). To probe whether the stimulatory effect of the Sov N-terminus supports Panx SUMOylation also in cells, we transfected OSCs with siRNAs targeting *sov*. The extent of Panx SUMOylation was unchanged in Sov-depleted cells (Figure S6E). This was different compared to HP1-depleted cells, where SUMOylation of Panx was elevated (Figure 6F). Considering that in both, Sov- and HP1-depleted cells, transposons were de-repressed to similar extents (Figure 2H), the unchanged SUMOylation of Panx in Sov-depleted cells might be the consequence of two opposing effects canceling each other out: increased recruitment of SFiNX to chromatin, yet decreased Panx SUMOylation efficiency in the absence of Sov. Taken together, our data reveal the elaborate molecular relationship between the Panx IDR and the Sov N-terminus (Figure S6F). The two-tiered interaction between SFiNX and Sov, which critically depends on SUMOylation of the Panx IDR by the core SUMO machinery, forms the molecular interface between the nuclear piRNA pathway and the heterochromatin machinery.

## DISCUSSION

With this work, we uncover the identity and control of the molecular interface between the nuclear piRNA pathway and the cellular heterochromatin machinery in *Drosophila* (Figure 7K). The core of this interface is a direct protein-protein interaction between the piRNA pathway-specific factor Panx and the ubiquitously expressed zinc finger Sov, which is required for general heterochromatin biology. Our central findings are that (1) Piwi-mediated stabilization of Panx at nascent transcripts on chromatin triggers SUMOylation of Panx and (2) that SUMOylation of Panx is critical for its interaction with Sov and thus for target silencing. We propose that induced SUMOylation of the SFiNX co-repressor subunit Panx on chromatin acts as a molecular switch to restrict functional interactions between the piRNA pathway and the general heterochromatin machinery to piRNA target sites.

We started our investigations at the level of SFiNX, the co-repressor complex consisting of Panx, Nxf2–Nxt1 and Ctp that acts at the interface of piRNA pathway and general heterochromatin machinery. Within SFiNX, we identified the N-terminal ∼200 amino acid region in Panx as the central and potentially only ‘silencing domain’. In several aspects, this region of Panx, termed IDR, resembles activating domains of transcriptional regulators (Sigler, 1988): it is intrinsically disordered, rich in prolines and, with a pK_a_ of 4.2, highly acidic. A common mode of how transcriptional regulators recruit co-activators or co-repressors is via ‘fuzzy’ interactions involving several weak hydrophobic interactions that in sum mediate specific binding (e.g. Tuttle et al., 2018). In the case of the Panx IDR, an amphipathic helix with central LxxLL motif binds directly to an alpha-helical domain (NTD) of the multi-zinc finger protein Sov. Remarkably, the NTD domain of Sov bears clear sequence similarity to a predicted folded domain in Med15, a Mediator subunit that binds to short motifs in transcriptional regulators. This might indicate that the Sov-NTD–LxxLL module was coopted from a binding module previously involved in transcriptional regulation. In isolation, the LxxLL–NTD interaction has an affinity of ∼1 micromolar. Given the low abundance of Panx and Sov in cells, this is potentially insufficient for meaningful binding in *vivo*. We find that an additional interaction between Panx and Sov, mediated by SUMOylation of the Panx IDR and two SIMs flanking the Sov NTD, is required for Panx function *in vivo*. Besides strengthening the Panx–Sov interaction, SUMOylation of Panx might also aid in exposing the LxxLL motif in the first place by disrupting intra-IDR interactions between the LxxLL motif and other hydrophobic sites. Consistent with this, strong consensus SUMOylation sites are located adjacent to the LxxLL motif and to a hydrophobic patch at the very N-terminus of the Panx IDR. Also, the 27 residue Panx peptide with LxxLL motif has a stronger silencing capacity (∼20-fold) than the full IDR with mutated SUMOylation sites (∼8-fold; Figure 1). These observations are consistent with the LxxLL motif being partially occluded in the context of the non-SUMOylated Panx IDR.

The concept of strengthening or coordinating protein-protein interactions through nearby SUMO-SIM pairs is a common cellular strategy to temporally and spatially constrain functional interactions between proteins (Kerscher, 2007). With this in mind, it is noteworthy that SUMOylated Panx isoforms are detectable only in the chromatin fraction, and that loss of Piwi leads to the absence of Panx SUMOylation. This suggests a simple model in which Piwi-mediated recruitment and/or stabilization of SFiNX on chromatin is mechanistically coupled to SUMOylation of Panx, thereby limiting a functional Panx-Sov interaction to piRNA target loci.

A central open question is how SUMOylation of Panx is molecularly restricted to piRNA target sites. This could be achieved through Piwi-engagement with a target transcript leading to the co-recruitment of substrate (SFiNX) and the SUMOylation machinery. Alternatively, the core SUMOylation machinery might be already present at chromatin and only the SFiNX substrate is recruited or stabilized on chromatin via Piwi. To understand this critical process, knowledge of the entire set of involved proteins will be required. Our data indicate that Su(var)2-10 is not required for Panx SUMOylation. As Panx harbors multiple SUMOylation consensus sites embedded in an acidic and intrinsically disordered polypeptide, it could be a direct Ubc9 substrate (Bernier-Villamor et al., 2002). Consistent with this, Panx is readily SUMOylated *in vitro* in an E3 independent manner. We note, however, that the involvement of an as yet unidentified E3 SUMO ligase cannot be ruled out at this stage. Based on our results, we currently favor a “substrate-to-enzyme” model in which Ubc9 is constitutively present on chromatin (e.g., at transcribed loci, as shown in Neyret-Kahn et al., 2013) and in which recruitment of Panx to chromatin, and thus to Ubc9, results in Panx SUMOylation. Consistent with this model, the isolated Panx IDR, whose silencing activity requires its SUMOylation sites, is a strong silencing domain independent of other SFiNX subunits or Piwi.

While recruitment of the Panx IDR to a reporter locus via DNA tethering results in strong silencing and heterochromatin formation, its recruitment to a nascent RNA, which mimics the actual *in vivo* situation, has no measurable effect on reporter expression. This is in stark contrast to the recruitment of full-length Panx, which is a very strong co-transcriptional silencer (Sienski et al., 2015; Yu et al., 2015; Batki et al., 2019). Such an on/off difference in silencing capacity suggests that the IDR, when recruited in isolation to a nascent RNA is not capable of establishing a functional interaction with Sov. We therefore propose that a key function of the structured Panx C-terminus, which interacts with Nxf2–Nxt1 and the dimerization unit Ctp, is to increase the residence time of the complex on chromatin, allowing for Panx SUMOylation and thus recruitment of Sov to the locus (Eastwood et al., 2021; Schnabl et al., 2021).

The molecular function of Sov in the establishment of heterochromatin at piRNA target loci is unclear. Its multiple C2H2 zinc fingers and reported interaction with HP1 suggest a role in recruitment and/or stabilization of HP1, a key factor in heterochromatin initiation and maintenance, on chromatin (Benner et al., 2019; Jankovics et al., 2018). However, experimental recruitment of HP1 to a nascent reporter transcript does not result in co-transcriptional silencing (Batki et al., 2019; Sienski et al., 2015), suggesting that Sov must have additional molecular activities. One of these may be related to Su(var)2-10, a SUMO E3 ligase of the PIAS family that is required for piRNA-mediated and general heterochromatin formation (Ninova et al., 2020a; Ninova et al., 2020b). Depletion of Su(var)2-10 in OSCs resulted in increased Panx SUMOylation. This was similar to depletion of HP1 and therefore places Su(var)2-10 downstream of the SUMOylation-dependent SFiNX-Sov interaction. We found Su(var)2-10 enriched in co-immunoprecipitation experiments with tagged Sov but not with tagged Panx (Figure S3A). E3 Ligases of the PIAS family have been implicated in protein group SUMOylation, where also accessible lysine residues outside of the Ψ-K-x-D/E consensus motif are SUMOylated (Jentsch and Psakhye, 2013; Li et al., 2020). This would create a multi-SUMO binding platform for diverse heterochromatin effector complexes harboring SIMs, such as the H3K9 methyltransferase Eggless–Windei (SetDB1–ATF7IP), histone deacetylases or the H3K4 demethylase Su(var)3-3 (Lsd1) (Ninova et al., 2020a). Accordingly, SUMOylation would play a dual role at piRNA target sites: As a regulatory switch, it gates the molecular interface between the piRNA pathway and Sov. And as an amplifier, with the critical role of the E3 SUMO ligase Su(var)2-10, it creates a “molecular glue” that recruits the various effector complexes whose combined action shuts down the locus through heterochromatin formation. Because Su(var)2-10 itself harbors SIMs, its initial recruitment, possibly via Sov, could then activate a feed-forward amplification loop. This model has conceptual parallels to double-stranded DNA damage repair, in which the initial trigger leads to recruitment of primary factors to the site of DNA damage and subsequently, a cascade of protein-group SUMOylation provides a binding platform for the various factors required for efficient DNA break repair (Jentsch and Psakhye, 2013).

## ACKNOWLEDGMENTS

We thank the VBCF core facilities (Protein Technologies, NGS, VDRC) for support and the IMBA Fly House for generating transgenic and CRISPR-edited fly lines. The GMI/IMBA/IMP Scientific Service units, especially the Mass Spectrometry unit (Karl Mechtler & team) provided outstanding support. We thank the Brennecke lab for help throughout this project, and Plamen Batalski for generating amino acid mapping scripts. Cristopher Lima and Ulrich Hohmann gave valuable feedback on the manuscript. The Brennecke lab is funded by the Austrian Academy of Sciences, the European Community (ERC-2015-CoG - 682181), and the Austrian Science Fund (F 4303 and W1207). X-ray diffraction studies were conducted at the Advanced Photon Source on the Northeastern Collaborative Access Team beamlines, supported by NIGMS grant P30 GM124165 and U.S. Department of Energy grant DE-AC02-06CH11357. The Eiger 16M detector on 24-ID-E beamline is funded by a NIH-ORIP HEI grant (S10OD021527). This work was supported in part by the Maloris Foundation (DJP). The Memorial Sloan Kettering Cancer Center structural biology core facility is supported by National Cancer Institute Core grant P30-CA008748. C. Yu and M. Gehre are supported by the VIP^2^ Post-Doc fellowship program as part of the EU Horizon 2020 research and innovation program (Marie Skłodowska-Curie grant No. 847548).

## AUTHOR CONTRIBUTIONS

J.W. undertook X-ray studies on the Sov NTD – Panx LxxLL peptide complex and ITC assays under supervision of D.J.P. V.I.A. and C.Y. performed all molecular biology experiments, except: J.S. identified the LxxLL motif and purified recombinant SFiNX complex, J.B., L.T., L.B. and P.D. performed fly experiments, M.G. performed H3K9me3 Cut&Run experiments and G.S. generated GRO-Seq data. C.Y., D.H. and C.Y. performed computational analyses, M.N. performed the phylogenetic analysis of the Panx LxxLL peptide, L.B. established *sov* RNAi lines and the Sov antibody, K.M. generated OSC cell lines. The project was supervised by J.B. and D.J.P. The paper was written by V.I.A, C.Y. and J.B. with input from all authors.

## MATERIALS AND METHODS

### Fly strains

All fly strains used in this study are listed in Supplementary Table 2 and are available from the VDRC (http://stockcenter.vdrc.at/control/main). Flies used for fertility scoring and ovary analysis were aged for 4 days at 25°C on apple juice agar with yeast paste before analysis. *panx* rescue strains were generated as previously described (Batki et al., 2019). For mutagenesis of the two SIMs in *sov* two pairs of sgRNAs (1+4 and 3+2) were cloned into pCFD4d (Addgene 83954) (Ge et al., 2016) as described (Port et al., 2014). A 1kb fragment of the N-terminus with modified SIM domains and guide target sites was synthesized (Genewiz) and amplified by PCR with two primer pairs to yield a shorter (750bp) and a longer (980bp) product. Both PCR products were mixed in equimolar amounts, denatured and reannealed (Dokshin et al., 2018). Two pCFD4d plasmids (each at 40ng/µl) expressing four different sgRNAs were co-injected with 100ng/µl Hsp70-Cas9 (Addgene 45945) (Gratz et al., 2013) and 100ng/µl of the hybrid dsDNA donor into white embryos. Flies containing the desired nucleotide changes were identified by PCR and subsequent Sanger sequencing. For the generation of the double K73E K80E (2KE) *sov* mutant two guides (Supplementary Table 3) were cloned into pDCC6 (Addgene 59985) and co-injected with an AltR HDR donor oligo (IDT) into white embryos as described (Gokcezade et al., 2014). F2 flies were screened by PCR and subsequent Sanger sequencing to identify those with the desired nucleotide changes. The boxB-GFP sensor was inserted into attP33 (Markstein et al., 2008) and recombined with the UASP-λN-HA-Panx transgene in attP40. The resulting boxB-GFP sensor UASP-λN-HA-Panx stock was combined with the *sov* or *white* sh-RNA transgenes (Ni et al., 2011) on the third chromosome, and crossed to the MTD-Gal4 driver line.

### OSC culture and siRNA transfection

OSCs were cultured as previously described (Niki et al., 2006; Saito et al., 2009). For plasmid DNA and siRNA transfections, Cell Line Nucleofector Kit V (Amaxa Biosystems) was used with program T-029. 5 × 10^6^ cells were used per transfection with 250 pmol siRNA duplexes (Supplementary Table 4).

### Generation of endogenously tagged OSC cell lines

Panx was N-terminally tagged at its endogenous locus with an HDR cassette consisting of a puromycin-resistance gene followed by a P2A cleavage site linked to FLAG-GFP. For Sov C-terminal tagging, the order of the puromycin-resistance gene and FLAG-GFP were reversed. The HDR cassette was flanked by ∼500-bp homology arms around the start codon and stop codon in *panx* and *sov* respectively and cloned into a pBluescriptII SK (+) backbone. 2500 ng of purified HDR PCR product and 1500 ng of guide RNA expression plasmid (Addgene 49330) containing the relevant guide RNAs (Supplementary Table 3) were transfected into 5 × 10^6^ OSCs. 24 h post transfection puromycin-containing medium (5 *µ*g/mL) was added to the cells and resistant clones were allowed to grow for 10 days. Individual clones were picked and analyzed by PCR, western blot and FACS for successful integration of the tagging cassette.

### Reporter tethering assay

The GFP reporter tethering assay in OSCs was carried out as described (Batki et al., 2019). In brief, CDS fragments of interest were cloned into the entry Gal4 tethering vector (Addgene 128013-128014) and the resulting plasmids were electroporated into OSCs, followed by treatment with puromycin to enrich for the transfected population. 4 days after transfection, cells were analyzed by flow cytometry on a FACS BD LSR Fortessa (BD Biosciences). Transfected cells were gated using mCherry expression and GFP fluorescence was measured for the population (10,000 cells per experiment). Data analysis was carried out in FlowJo (FlowJo, LLC).

### Immunofluorescence staining of ovaries

5-10 ovary pairs were dissected into ice-cold PBS and fixed in immunofluorescence Fixing Buffer (4 % formaldehyde, 0.3 % Triton X-100, PBS) for 20 minutes at room temperature with rotation. Fixed ovaries were then washed 3 times with PBX (0.3 % Triton X-100, 1x PBS), and blocked in BBX (0.1 % BSA, 0.3 % Triton X-100, 1x PBS) for 30 min at room temperature. The samples were incubated with primary antibody in BBX overnight at 4°C with rotation, followed by 3 in PBX and an overnight incubation with secondary antibody in BBX at 4°C with rotation. After rinsing with PBX, samples were stained for 5 min with 0.1 *µ*g.mL^-1^ DAPI and washed 3 times with PBX. Ovaries were resuspended in ∼50 *µ*l DAKO Fluorescence mounting medium and imaged on a Zeiss LSM-880 Axio Imager confocal microscope. For GFP reporter imaging ovaries were washed with PBX after fixation and stained with 0.1 *µ*g.mL^-1^ DAPI followed by 3 washed with PBX. Ovaries were resuspended in ∼20 *µ*L VectaShield mounting medium, imaged on a Zeiss LSM-880 Axio Imager confocal microscope equipped with AiryScan detector and the resulting images processed using FIJI/ImageJ (Schindelin et al., 2012). Antibodies are listed in Supplementary Table 5.

### Generation of Sov and Su(var)2-10 antibodies

Purified His6-tagged Sov (aa 90-297) was used to generate the anti-Sov monoclonal antibody used for western blot and immunofluorescence. For Su(var)2-10, we raised a monoclonal antibody against the His6-tagged region corresponding to amino acids 2–514. Su(var)2-10 and Sov mouse monoclonal antibodies were generated by the Max F. Perutz Laboratories Monoclonal Antibody Facility.

### Whole cell extract preparation and subcellular fractionation

For whole cell extracts (WCE) cells were washed once with PBS and resuspended in ice-cold WCE buffer (10 mM Tris-HCl, 1% NP-40, 2 mM MgCl_2_, benzonase, cOmplete Protease Inhibitor Cocktail (Roche), 25 nM NEM where relevant) for 15 min on ice. After protein concentration measurement, the lysates were boiled with 1x final concentration SDS-PAGE loading buffer at 95 °C for 3 min. For subcellular fractionation the cell pellet was resuspended in CSK buffer (10 mM HEPES-KOH, pH 7.3, 300 mM sucrose, 0.5 % Triton X-100, 100 mM NaCl, 3 mM MgCl_2_, 25 nM N-ethylmaleimide (NEM), cOmplete Protease Inhibitor Cocktail (Roche)) for 4 minutes on ice followed by centrifugation at 2500 × g for 5 min at 4 °C. The supernatant was transferred to a new tube and further centrifuged at 20,000 × g for 15 min at 4 °C to remove cellular debris and kept as the soluble fraction. The pellet was washed once in CSK buffer, resuspended in WCE buffer, and then boiled in 1x final concentration SDS-PAGE loading buffer at 95 °C for 3min. All fractions were prepared in the same volume as the WCE samples.

### Western blotting

Proteins were separated by SDS polyacrylamide gel electrophoresis and transferred to 0.2 *µ*m nitrocellulose membrane (Bio-Rad). Protein transfer and equal loading were checked by staining with Ponceau S. The membrane was blocked with 5 % non-fat milk in PBX (PBS, 0.05 % Triton X-100). Following blocking, the membranes were incubated with primary antibodies overnight at 4 °C or for 1 hour at room temperature. After primary antibody incubation the membrane was washed 3 times with PBX for 5 min followed by incubation with HRP-conjugated secondary antibodies in 5 % milk in PBX for 1 hour at room temperature. The membrane was then washed 3 times for 5 minutes with PBX, incubated with Clarity Western ECL Blotting Substrate (Bio-Rad) and imaged with a ChemiDoc MP imager (Bio-Rad). Antibodies are listed in Supplementary Table 5.

### Protein co-immunoprecipitation from S2 cell lysates

8 × 10^6^ S2 cells were transfected with 2 *µ*g plasmids encoding FLAG- and GFP-tagged fragments of interest from Panx and Sov respectively using Cell Line Nucleofector Kit V (Amaxa Biosystems) with program G-030. 2 days after transfection, cells were collected, washed once with cold PBS and resuspended in S2 lysis buffer (50 mM Tris-HCl pH 7.5, 150 mM NaCl, 0.5 % Triton X-100, 10 % glycerol, 1 mM DTT, cOmplete Protease Inhibitor Cocktail (Roche)). After incubation for 30 min on ice the cell lysate was centrifuged (20,000 × g for 15 min at 4 °C) and the supernatant was collected. Magnetic agarose GFP-Trap beads (ChromoTek) were incubated with the lysate for 2 hours at 4 °C with nutation. Subsequently, the beads were washed 3 x for 10 min with S2 lysis buffer, boiled in 2x SDS-PAGE loading buffer for 5 minutes at 95 °C and the eluate analyzed by western blotting.

### co-immunoprecipitation of SUMOylated proteins with denaturing wash step from fly ovaries

FLAG-GFP-Smt3 expressing and wild type control flies were dissected and 200 *µ*L of ovaries were washed once with ice-cold PBS and dounced (40 times) in 1 mL ovary lysis buffer (OLB) (50 mM Tris-HCl pH 8, 150 mM NaCl, 0.25 % Triton X-100, 0.3 % NP-40, 10 % glycerol, 2 mM MgCl_2_, 25 mM N-ethylmaleimide (NEM), cOmplete Protease Inhibitor Cocktail (Roche)). The ovary lysate was incubated for 30 min at 4 °C with nutation and centrifuged for 5 min at 20,000 × g at 4 °C. The supernatant was kept on ice and the pellet was resuspended in 200 *µ*L OLB with 500 mM NaCl and sonicated for 10 min at high setting with 30 s / 30 s duty cycle on a Bioruptor sonicator (Diagenode) and centrifuged for 5 min at 20,000 × g at 4 °C. The supernatant from this step was combined with the supernatant from the previous step and centrifuged for 5 min at 20,000 × g at 4 °C. The resulting supernatant was pre-cleared by incubation with Sepharose beads for 30 min at 4 °C and centrifuged for 5 min at 20,000 × g at 4 °C. The supernatant was extracted with a syringe to bypass the lipid layer on top and mixed with OLB pre-equilibrated magnetic agarose GFP-Trap beads (ChromoTek) followed by overnight incubation at 4 °C. The beads were washed once with OLB for min 10 min at 4 °C, once with RIPA buffer (10 mM Tris-HCl pH 8 150 mM NaCl, 0.5 mM EDTA, 0.1 % SDS, 1 % NP-40, 1% sodium deoxycholate) for min 10 min at 4 °C followed by a single wash with high salt wash buffer (10 mM Tris-HCl pH 8, 1M NaCl, 0.1 % NP-40) for min 10 min at 4 °C and then 2 washes with SDS urea wash buffer (10 mM Tris-HCl pH 8, 1 % SDS, 8 M urea) for min 10 min at room temperature. Beads were further processed for downstream mass spectrometry analysis and a small aliquot boiled in 2x SDS-PAGE loading buffer at 95 °C for 5 min and analyzed by SDS-PAGE followed by silver staining (Pierce Silver stain Kit, #24612).

### Recombinant protein pulldown and co-immunoprecipitation from OSC extracts

5 × 10^8^ OSCs were washed in PBS and resuspended in ice-cold CSK buffer (10 mM HEPES-KOH pH 7.3, 300 mM sucrose, 0.5 % Triton X-100, 100 mM NaCl, 3 mM MgCl_2_, 25 nM N-ethylmaleimide (NEM), cOmplete Protease Inhibitor Cocktail (Roche)) and incubated for 5 min on ice followed by centrifugation at 2500 × g for 5 min at 4 °C. The pellet was resuspended in RIPA buffer (50 mM HEPES-KOH, pH 7.3, 200 mM KCl, 3.2 mM MgCl_2_, 0.25 % Triton X-100, 0.25 % NP-40, 0.1 % sodium deoxycholate, 10 % glycerol, benzonase, 25 mM NEM) and incubated for 1 hour at 4 °C with nutation. The lysate was centrifuged at 18,000 × g for 10 min at 4 °C and supernatant was collected for either pulldowns or co-immunoprecipitation. For recombinant protein pulldown, GFP-tagged variants of the Sov NTD were immobilized on magnetic agarose GFP-Trap beads (ChromoTek) by incubation in PBS with 0.1 % Triton X-100 for 3 hours at 4 °C. The GFP-fusion pre-coupled beads were incubated with OSC extract for 3 hours at 4 °C with nutation. The beads were washed 3 x for 10 min at 4 °C with RIPA buffer and associated proteins were eluted by boiling in 2x SDS-PAGE loading buffer for 5 min at 95 °C. The eluate was analyzed by western blotting. For co-immunoprecipitation, the extract prepared from OSC lines expressing endogenously tagged proteins of interest was incubated with magnetic GFP-Trap agarose (Chromotek) for 3h at 4°C. The beads were washed 3 x with RIPA buffer for 10 min at 4 °C, followed by 3 x washes with non-detergent buffer (20 mM HEPES pH 7.4, 137 mM NaCl). 80% of the washed beads were resuspend in 100mM ammonium bicarbonate and used for mass spectrometry analysis. 20% of the beads-associated proteins were eluted in 2x SDS-PAGE loading buffer by incubating at 95°C for 5 min. The eluate was analyzed by silver staining.

### Peptide pulldowns

Peptides corresponding to Panx aa 82-108 (WT LxxLL and mutant NxxQQ) were chemically synthesized with an aminohexanoate linked N-terminal biotin moiety and a C-terminal amide blocking group. Peptides were precoupled to streptavidin magnetic beads (Pierce) in PBS with 0.1 % Triton X-100 for 3 hours at 4 °C. For nuclear extract pulldown, 5 × 10^8^ OSCs were resuspended in hypotonic buffer (10 mM Tris-HCl pH 7.5, 2 mM MgCl_2_, 3 mM CaCl_2_, 1 mM DTT, cOmplete EDTA-free protease inhibitor cocktail (Roche)) for 10 min at 4 °C followed by centrifugation (500 × g for 5 min at 4 °C). The pellet was resuspended in hypotonic lysis buffer (10 mM Tris-HCl pH 7.5, 2 mM MgCl_2_, 3 mM CaCl_2_, 0.5 % Igepal CA-630, 10 % glycerol, 1 mM DTT, cOmplete EDTA-free protease inhibitor cocktail (Roche)) and incubated for 20 min at 4 °C. After centrifugation, the resulting nuclear pellet was resuspended in nuclear lysis buffer (20 mM HEPES-KOH, pH 7.9, 150 mM NaCl, 0.3 % Triton X-100, 0.25 % NP-40, 10 % glycerol, 1 mM DTT, cOmplete EDTA-free protease inhibitor cocktail (Roche)) and incubated with nutation for 30 min at 4 °C. For Sov peptide mapping, the nuclear extract was sonicated for 15 min at high intensity (30s/30s duty cycle); Bioruptor sonicator (Diagenode)). After lysate centrifugation (20,000 × g for 15 min at 4 °C), the supernatant was collected and incubated for 3 hours at 4 °C with the peptide pre-coupled beads. Beads were washed 3 x in nuclear lysis buffer, followed by detergent-free wash step and sent for mass spectrometry analysis. For *in vitro* binding assays, peptides were coupled to magnetic streptavidin beads (Pierce) and incubated with recombinant Sov fragments in peptide binding buffer (50 mM Tris-HCl pH 7.5, 150 mM NaCl, 10 % glycerol, 0.5 % Triton X-100) for 3 hours at 4 °C followed by 3 washes for 10 min in the same buffer. Beads were boiled in 2x SDS-PAGE loading buffer for 5 min at 95 °C and the eluate was analyzed by SDS-PAGE followed by Coomassie staining.

### Recombinant Sov fragment purification for pulldown assays

Sov fragments of interest were cloned in a pET15b bacterial expression vector carrying a C-terminal GFP His6 tag. Transformed *E. coli* strain BL21(DE3) were grown at 37 °C until OD_600_ reached 0.8 then the culture was cooled down to 18 °C and induced with 0.1 mM IPTG for 18 hours. Cell pellets were resuspended in NTD lysis buffer (50 mM sodium phosphate, pH 8, 200 mM NaCl, 0.1 % Triton X-100, 10% glycerol, 10 mM imidazole, 5 mM β-mercaptoethanol, 1 mM PMSF) and passed twice through a French press followed by ultracentrifugation at 100,000 × g for 1 hour at 4 °C. The supernatant was passed through a 5 ml HisTrap HP column (Cytiva, #17524801) and bound protein was eluted in a gradient with NTD lysis buffer containing 500 mM imidazole and no Triton X-100. Fractions containing the fragment of interest were pooled and diluted to a final concentration of 50 mM NaCl and ran through HiTrap Q anion exchange column (Cytiva, #17115401) followed by elution in 50 mM Tris pH 8 with a linear 50-500 mM NaCl gradient. Fractions of interest were pooled and concentrated on a 15 kDa MWCO spin concentrator (Sartorius, #VS2001) and were then loaded onto a Superdex 75 gel filtration column (GE Healthcare, #28-9893-33) (equilibrated in 50 mM Tris pH 8 and 150 mM NaCl). Fractions containing the protein of interest were pooled, aliquoted and stored at −80 °C.

### Recombinant IDR and SFiNX complex purification

Panx IDR (aa 1-195) was cloned in a pET21a bacterial expression vector with a C-terminal 3xHA-His10 tag and transformed into *E. coli* strain BL21(DE3). Bacteria were grown at 37 °C until OD_600_ reached 0.8 and then induced by 1 mM IPTG for 2 hours at 37 °C. The bacterial pellet was resuspended in IDR lysis buffer (50 mM sodium phosphate, pH 8, 300 mM NaCl, 10 mM imidazole) and freeze-thawed with the addition of 5 mM β-mercaptoethanol and 1 mM phenylmethylsulfonyl fluoride. The cell suspension was passed twice through a French press and then ultracentrifuged at 100,000 × g for 1 hour at 4 °C. The supernatant was passed through a 5 ml HisTrap HP column (Cytiva, #17524801) and bound protein was eluted in a gradient with IDR lysis buffer containing 500 mM imidazole. Fractions that contained the IDR were pooled and diluted to a final concentration of 50 mM NaCl and passed through HiTrap Q anion exchange column (Cytiva, #17115401) followed by elution in 50 mM Tris pH 7.5 with a linear 50-500 mM NaCl gradient. Fractions containing the protein of interest were pooled and concentrated on a 5 kDa MWCO spin concentrator (Cytiva, #28-9323-59). Urea was added to a final concentration of 8M followed by overnight dialysis against transport buffer (20 mM HEPES, pH 7.3, 110 mM potassium acetate, 2 mM magnesium acetate, 1 mM EGTA, 1 mM DTT). The protein was then loaded onto a transport buffer equilibrated Superdex 75 gel filtration column (GE Healthcare, #28-9893-33) and fractions containing the protein of interest were pooled, aliquoted and stored at −80 °C. Full length SFiNX was purified as described (Schnabl et al., 2021) with the exception that size exclusion chromatography was performed with a HiLoad 16/60 Superose 6 column (Cytiva, #29323952).

### RNA preparation and RT-qPCR

5-10 million OSCs or 5–10 pairs of ovaries were collected, and total RNA was isolated using NucleoSpin RNAXS kit (Macherey-Nagel) according to the manufacturer’s instructions. Complementary DNA was prepared using 1 *µ*g total RNA and random hexamer oligonucleotides with SuperScript IV Reverse Transcriptase (Invitrogen). Primers used for qPCR analysis are listed in Supplementary Table 6.

### RNA-seq and RNA-seq analysis

For RNA-seq libraries, total RNA was isolated with TRIzol reagent, and poly(A)+ RNA was enriched with Dynabeads Oligo(dT)25 (Thermo Fisher, 61002) according to the user manual. cDNA was prepared with NEBNext Ultra II RNA First and Second Strand Synthesis Module, followed by library preparation with NEBNext Ultra II DNA Library Prep Kit Illumina (NEB). The library was sequenced on HiSeqV4 (Illumina) in SR50 mode. RNA-seq analysis and differential gene expression analysis were carried out as described (Batki et al., 2019).

### ChIP-seq

ChIP was performed according to (Lee et al., 2006) with modifications, except for H3K9me3 ChIP after Sov-tethering, which was performed using ultra-low-input micrococcal nuclease-based native ChIP (ULI-NChIP) according to (Brind’Amour et al., 2015). For ChIP after pre-extracting cells, OSCs were first treated on ice with CSK buffer (10 mM HEPES-KOH, pH 7.3, 300 mM sucrose, 0.5 % Triton X-100, 100 mM NaCl, 3 mM MgCl_2_, 25 nM N-ethylmaleimide (NEM), cOmplete Protease Inhibitor Cocktail (Roche)) for 5 min before crosslinking; otherwise, OSCs were directly crosslinked with 1% formaldehyde for 10min at RT, the reaction was quenched with glycine (final concentration of 125mM) and cells were washed twice with cold 1x PBS. Chromatin was prepared using lysis buffer (50 mM Tris-HCl, pH 8.0, 2 mM EDTA, 0.5% NP-40, 10% Glycerol,1× protease inhibitors), followed by sonication in sonication buffer (20 mM Tris-HCl, pH 8.0, 150 mM NaCl, 2 mM EDTA, 0.2% SDS, 1× protease inhibitors) with Covaris E220 sonicator for 6 min (Duty Factor 5.0, Peak Incident Power 140, Cycles per Burst 200). Protein G and Protein A Dynabeads (1:1 mixed) were blocked with 1mg/ml denatured yeast tRNA (Sigma-Aldrich, R5636) and 1mg/ml of BSA (NEB, B9000S) for 2h at 4 °C, and then anti-GFP antibody (Thermo Scientific, A-111222) or anti H3K9me3 antibody (Active Motif, 39161) were coupled to the blocked Dynabeads for 2h at 4 °C. The antibody-coupled beads were then added to the sheared chromatin and incubated overnight at 4 °C. Beads were washed with wash buffer 1 (20 mM Tris-HCl, pH 8.0, 500 mM NaCl, 2 mM EDTA, 0.5% NP-40, 0.1% SDS) and wash buffer 2 (10 mM Tris-HCl, pH 8.0, 250 mM LiCl, 1 mM EDTA, 0.5% NP-40, 0.5% Na-Deoxycholate), followed by elution with elution buffer (0.1M NaHCO_3_, 1% SDS). Eluates and inputs were de-crosslinked at 65 °C overnight. Following RNase A and proteinase K treatment, DNA was purified via Phenol/Chloroform extraction. ChIP-seq libraries were prepared using NEBNext Ultra DNA Library Prep Kit Illumina (NEB) and sequenced on HiSeqV4 (Illumina) in SR50 mode.

### ChIP-seq analysis

ChIP-seq analysis was carried out as described (Batki et al., 2019). In brief, after removal of the adaptors, sequencing reads with a minimal length of 18 nt were mapped to the *D. melanogaster* genome (dm6) using Bowtie (Langmead, B., et al. 2009, PMID: 19261174) (release v.1.2.2), with zero mismatch allowed for genome wide analysis. For the TE-consensus analysis, reads were mapped allowing ‘0’ mismatches and multi-mapping only within a single transposon. BigWig files were generated using Homer (Heinz et al., 2010) and UCSC BigWig tools (Kent et al., 2010). Heatmaps were generated with Deeptools (Ramirez et al., 2016) using BigWig files with uniquely mapped reads, and meta profiles were generated with ngs.plot (Shen et al., 2014) using bam files with uniquely mapped reads. The Piwi-regulated TEs were used as in (Sienski et al., 2015). All sequenced libraries are listed in Supplementary Table 7 and are accessible via GEO (accession #GSE173237).

### Mass spectrometry

Mass spectrometry was carried out as described (Batki et al., 2019).

### *In vitro* SUMOylation assay

Coding sequences for Aos1, Uba2, Lwr and Smt3 were cloned from *Drosophila* cDNA. E1, E2 and Smt3 protein expression and purification was carried out as described in (Flotho et al., 2012). *In vitro* SUMOylation reactions were assembled in SUMOylation assay buffer (20 mM HEPES, pH 7.3, 110 mM potassium acetate, 2 mM magnesium acetate, 1 mM EGTA, 1 mM DTT, 0.05 % Tween-20, 20 *µ*g/mL BSA) with 150 nM His-Aos1/Uba2, 500 nM Lwr, 2 *µ*M Smt3 (unless stated otherwise). Additional proteins added to the reaction mix were dialyzed overnight against transport buffer (20 mM HEPES, pH 7.3, 110 mM potassium acetate, 2 mM magnesium acetate, 1 mM EGTA, 1 mM DTT) to avoid dilution effects on reaction kinetics. 20 *µ*L reactions were assembled on ice and initiated by the addition of 5 mM ATP, incubated for 1 hour at 30 °C and terminated with 20 *µ*L 2x SDS-PAGE loading buffer followed by western blot analysis.

### Cut&Run

Cut&Run was performed according to (Skene et al., 2018) with minor modifications. In brief, OSCs were harvested and 500.000 OSCs were washed three times and coupled to 10 *µ*l of activated Concanavalin A-coated magnetic beads (Polysciences, #86057-3) per sample. Cells were gently lysed using Dig-wash buffer (20 mM HEPES pH 7.3, 150 mM NaCl, 0.5 mM spermidine, 0.01% digitonin, cOmplete Protease Inhibitor (Roche)) and incubated with 0.5 *µ*g of H3K9me3 (Active motif, #39161) or IgG (CST, #2729S) antibody at 4°C overnight on a nutator. Cells were washed and resuspended in Dig-wash buffer containing 700 ng/ml pAG-MNase (in-house production by Molecular Biology Facility) and incubated for 2 hours at 4°C on a nutator. After washing, cells were resuspended in Dig-wash buffer containing 2 mM CaCl_2_ to activate pAG-MNase. MNase activity was quenched by the addition of 2x STOP Buffer (340 mM NaCl, 20 mM EDTA, 4 mM EGTA, 0.02% Digitonin, 50 μg/ml RNase A, 50 μg/ml Glycogen). DNA fragments were released into solution by incubating samples at 37°C mixing at 500 rpm for 10 minutes. Insoluble material was pelleted by centrifugation and 0.1% SDS and 0.2 μg/μl Proteinase K were added to the supernatant followed by incubation for 1 hour at 55°C. DNA was purified using a DNA purification kit (in house) and libraries were prepared following the manufacturer’s instructions with NEBNext Ultra II DNA Library Prep Kit for Illumina using a shortened elongation time of 15 seconds during PCR amplification. Sequencing was performed on a NextSeq550 using 75 bp single-end mode.

### Cut&Run Data Analysis

Cut&Run sequencing reads were aligned to the *D. melanogaster* reference genome (dm6 assembly) using Bowtie2 (Galaxy v. 2.3.4.3, Langmead et al. 2012) with zero mismatches allowed. Only non-duplicated, uniquely mapped reads were retained for further analysis. Plots to visualize the distribution of H3K9me3 density around Piwi-regulated transposon insertion sites (Sienski et al., 2015) were generated using ngs.plot. The standard error of mean (SEM) across the regions was calculated and shown as a semi-transparent shade around the mean curve. (v.2.61) (Shen et al., 2014). For transposon consensus analysis, genome mapping reads longer than 23 nt were mapped to transposon consensus sequences using STAR (Dobin et al., 2013) (v.2.5.2b; settings: --outSAMmode NoQS -- readFilesCommand cat --alignEndsType Local --twopassMode Basic --outReadsUnmapped Fastx -- outMultimapperOrder Random --outSAMtype SAM --outFilterMultimapNmax 1000 -- winAnchorMultimapNmax 2000 --outFilterMismatchNmax 3 --seedSearchStartLmax 30 -- outFilterType BySJout --alignSJoverhangMin 15 --alignSJDBoverhangMin 1). Multiple mappings were only allowed within one transposon and read counts were divided equally to the mapping positions. For plotting, read counts were normalized to 10 million sequenced reads, converted to bedgraph tracks using Bedtools (v.2.27.1) (Quinlan and Hall, 2010) and plotted in RStudio as a smoothed conditional means function using the loess method (ggplot2 version 3.3.0); semi-transparent shade depict data points without smoothing.

### GRO-Seq

GRO-Seq was performed as described in (Sienski et al., 2012).

### Protein expression and purification for crystallography

*D. melanogaster* Panoramix (residues 83-109) and Sov (residues 14-90) were cloned into a modified RSFduet-1 vector (Novagen) with an N-terminal His_6_-SUMO tag on Panx and no tag on Sov. Panx and Sov were co-expressed in *E. coli* strain BL21(DE3) RIL (Stratagene). The cells were grown at 37°C until OD_600_ reached 0.8, then the media was cooled to 16°C and IPTG was added to a final concentration of 0.35 mM to induce protein expression overnight at 16°C. The cells were harvested by centrifugation at 4°C and disrupted by sonication in Binding buffer (20 mM Tris-HCl pH 8.0, 500 mM NaCl, 20 mM imidazole) supplemented with 1 mM PMSF (phenylmethylsulfonyl fluoride) and 3 mM β-mercaptoethanol. After centrifugation, the supernatant containing complexes of Panx and Sov was loaded onto 5 ml HisTrap Fastflow column (GE Healthcare). After extensive washing with Binding buffer, the complex was eluted with Binding buffer supplemented with 500 mM imidazole. The His_6_-SUMO tag was removed by Ulp1 protease digestion during dialysis against Binding buffer and separated by reloading onto HisTrap column. The flow-through fraction was further purified by HiTrap Q FF column and Superdex 75 16/60 column (GE Healthcare). The pooled fractions were concentrated to 20 mg/ml in crystallization buffer (20 mM Tris-HCl pH 7.5, 200 mM NaCl, 1 mM DTT). For the seleno-methionine (SeMet) derivative protein, the cells were grown in M9 medium supplemented with lysine, threonine, phenylalanine, leucine, isoleucine, valine, and Se-methionine and purified as described above.

### Crystallization, data collection and structure determination

Crystals of native and SeMet derivative Panx–Sov complex were grown from the same solutions containing 0.1 M CHES pH 9.5, 30% (w/v) PEG 3000 using the sitting drop vapor diffusion method at 20°C. For data collection, the crystals were flash frozen (100 K) and collected on NE-CAT beam lines 24ID-C and 24ID-E at the Advanced Photo Source (APS), Argonne National Laboratory. The diffraction data were processed with XDS (Kabsch, 2010) and iMosflm (Battye et al., 2011). The structure of the Panx-Sov complex was solved by the single-wavelength anomalous diffraction (SAD) method using PHENIX (Adams et al., 2002). The automatic model building was carried out using the program PHENIX AutoBuild (Adams et al., 2002). The resulting model was completed manually using COOT (Emsley et al., 2010) and PHENIX refinement (Adams et al., 2002). The statistics of the diffraction and refinement data are summarized in Supplementary Table 8. Molecular graphics were generated with the PyMOL program (https://pymol.org/2/) and UCSF Chimera X (Goddard et al., 2018).

### Isothermal Titration Calorimetry (ITC)

His-SUMO tagged Panx peptides and C-terminal His tagged Sov protein were purified separately in the same buffer (20 mM Tris pH7.5, 150 mM NaCl, 2 mM β-mercaptoethanol). The titrations were performed on a MicroCal ITC200 calorimeter at 20°C. The exothermic heat of reaction was measured by 20 sequential injections of 0.72 mM His-SUMO tagged Panx peptides into 30 *µ*M Sov protein solutions with 120 s interval spacing. The data was fitted using the program Origin with ‘one set of sites’ model.

### Conservation analysis of the Panx LxxLL motif

Panx orthologous proteins from the melanogaster species group (taxid 32346; 19 species) were obtained from the NCBI nr protein database and aligned using mafft v7 with default settings. A segment corresponding to aminoacids in 82-108 Dmel panx was extracted, visualized using clustalx and used to derive a sequence logo using ggseqlogo (v0.1 in R3.6.2).

### Data and code availability statement

Coordinates and structure factors of Sov NTD in complex with the Panx LxxLL peptide were deposited in the Protein Data Bank (PDB; accession 7MKK. Sequencing data sets were deposited in the NCBI GEO archive (accession GSE173237). The proteomics data were deposited in the ProteomeXchange Consortium via the PRIDE partner repository (data set PXD025437). All custom code not referenced in the methods is available upon request.

## SUPPLEMENTAL FIGURE TITLES AND LEGENDS

**Figure S1 (related to Figure 1).**
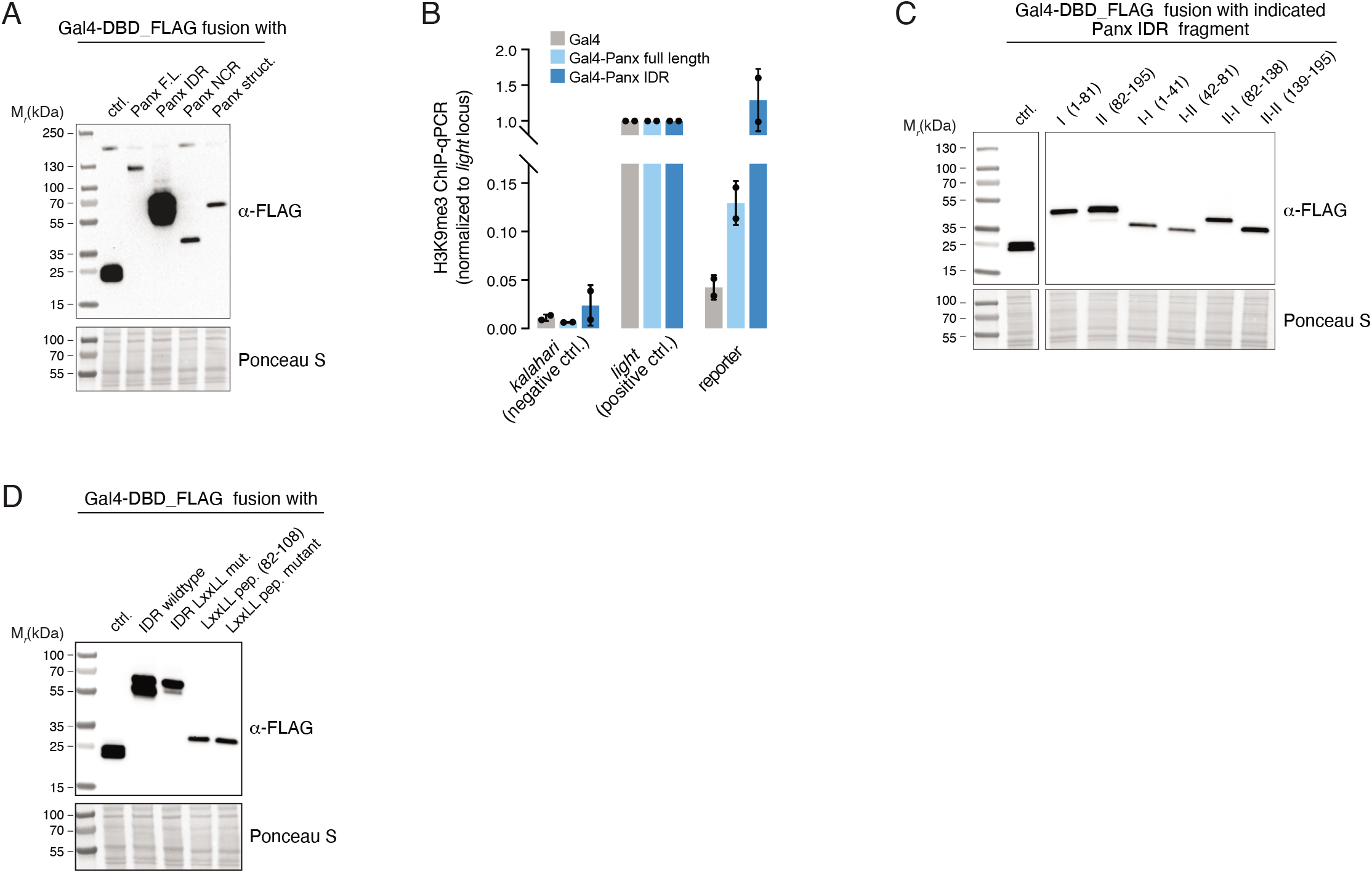
**A,** Western blot analysis showing expression levels of indicated Gal4-DBD fusion proteins following transient transfection in OSCs (Ponceau S levels indicate protein loading; related to Fig. 1B). **B,** H3K9me3 levels (normalized to heterochromatic *light* locus) at the reporter locus (amplicon indicated in Fig. 1A) determined by ChIP-qPCR from OSCs expressing indicated Gal4-DBD fusion proteins (n = 2 biological replicates; error bars: St. dev.). **C,** Western blot analysis showing expression levels of Gal4-DBD fusion proteins with indicated Panx IDR fragments (amino acid boundaries indicated). Ponceau S levels indicate protein loading; related to Fig. 1D. **D**, Western blot analysis showing expression levels of Gal4-DBD fusions with indicated WT and LxxLL mutant Panx fragments. Ponceau S levels indicate protein loading; related to Fig. 1E.

**Figure S2 (related to Figure 2).**
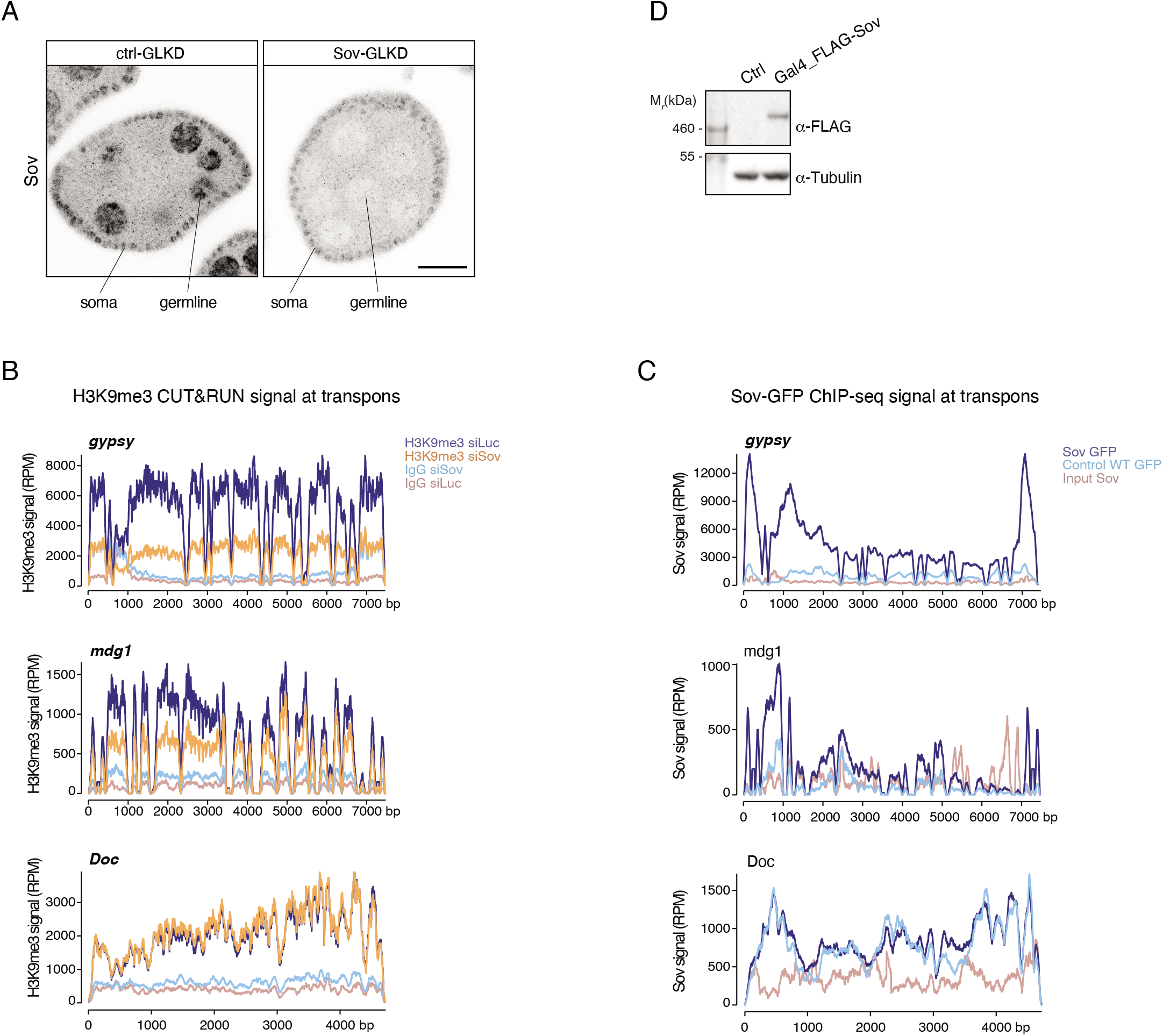
**A,** Confocal image of egg chambers with indicated germline-specific knockdown (GLKD) stained for Sov (greyscale; scale bar: 20 μm). **B**, H3K9me3 Cut&Run signal from OSCs with indicated knockdowns at indicated transposons (The *Doc* retroelement serves as a control transposon as it is not targeted by the piRNA pathway in OSCs). **C**, Sov-GFP ChIP-Seq signal at indicated transposons from OSCs expressing Sov-GFP or not (control) (The *Doc* retroelement serves as a control transposon as it is not targeted by the piRNA pathway in OSCs). **D**, Western blot analysis showing expression of Gal4-DBD-FLAG-Sov following plasmid transfection in the OSC reporter cell line (related to Figure 2F, G).

**Figure S3 (related to Figure 4).**
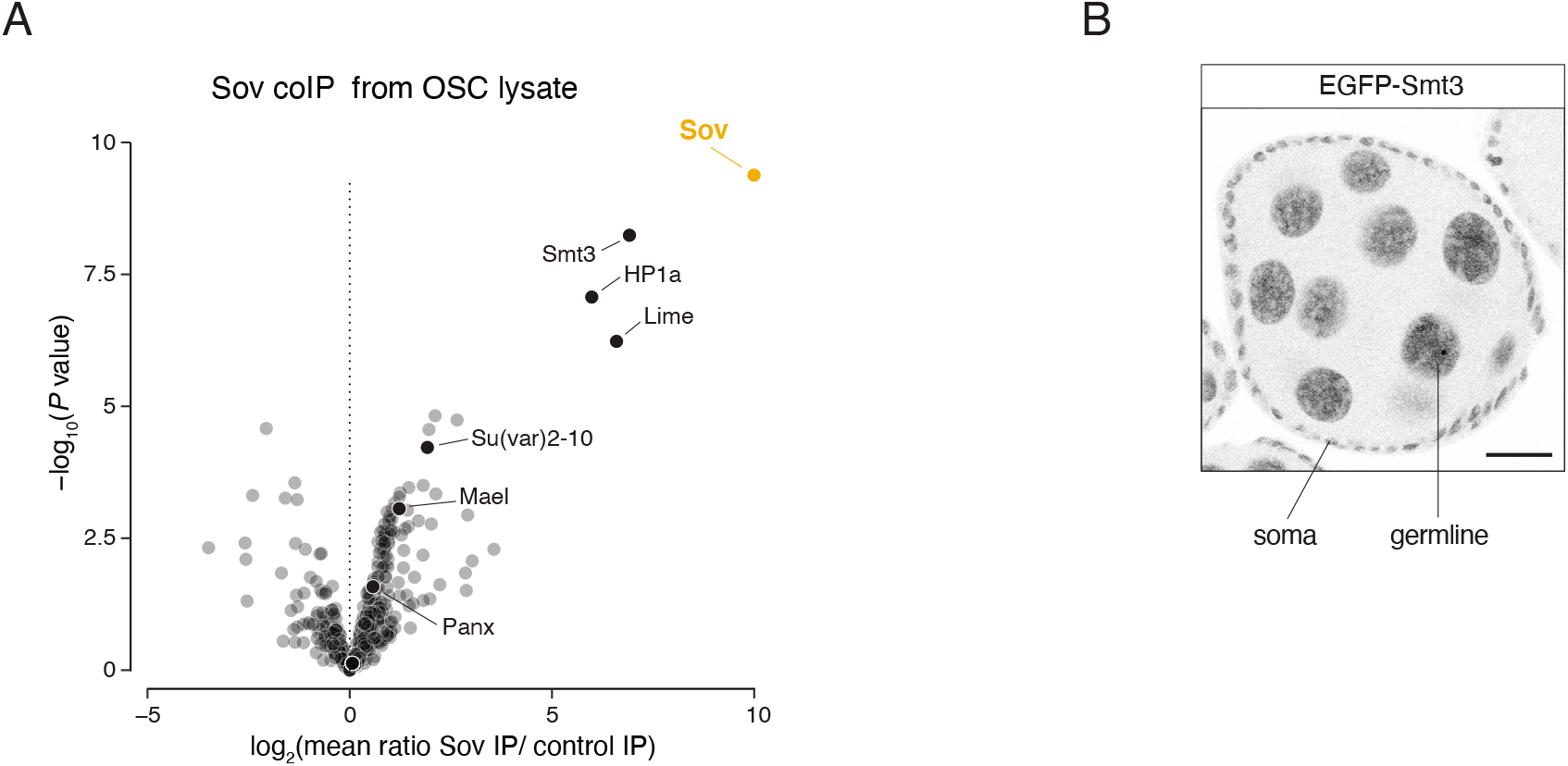
**A**, Volcano plot showing fold enrichment of proteins determined by quantitative mass spectrometry in GFP-FLAG-Sov co-immunoprecipitates versus control (*n* = 3 biological replicates; p-values corrected for multiple testing). **B**, Confocal image of egg chamber expressing GFP-Smt3 (greyscale) under the *smt3* regulatory control elements (scale bar: 20 μm).

**Figure S4 (related to Figure 5).**
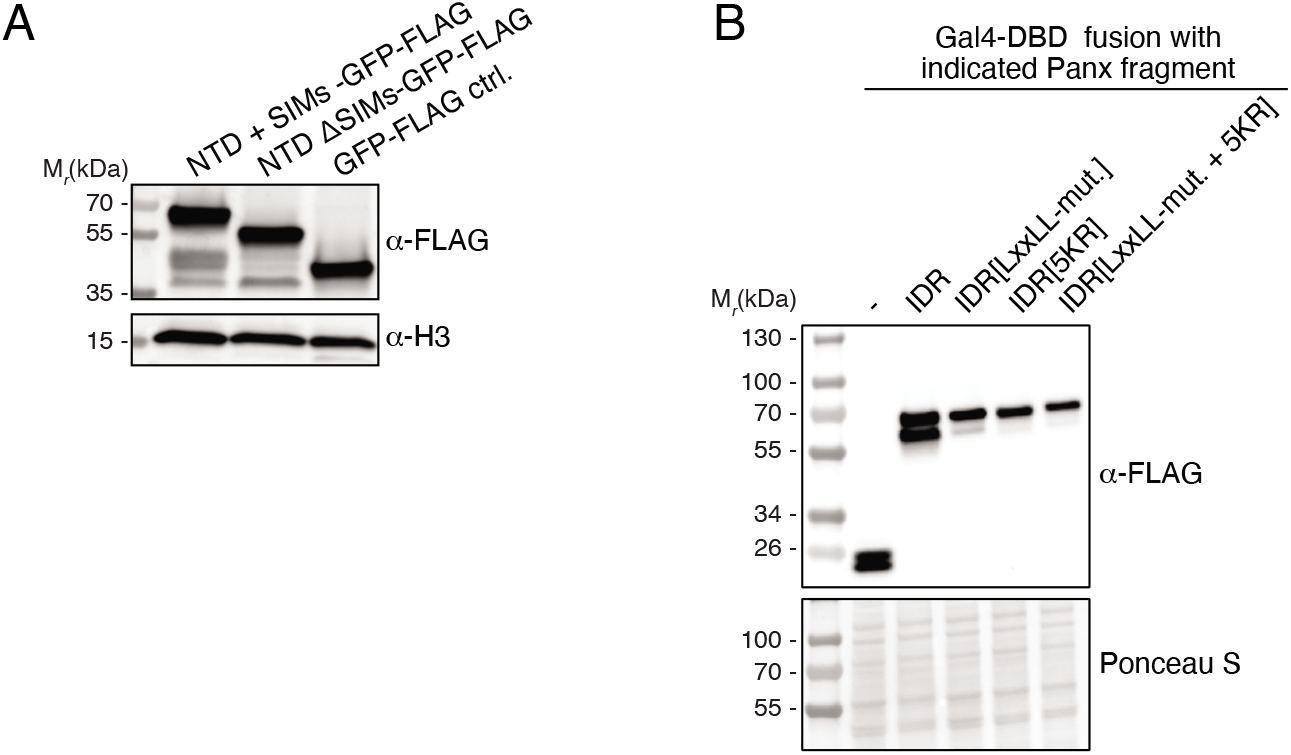
**A**, Western blot analysis showing expression of indicated GFP-FLAG tagged Sov NTD variants following transient transfection in OSCs (related to Figure 5A). **B**, Western blot analysis showing expression levels of Gal4-DBD-FLAG fusions with indicated Panx IDR variants following transient transfection in the OSC reporter line. Staining with Ponceau S serves as loading control (related to Figure 5B).

**Figure S5 (related to Figure 6).**
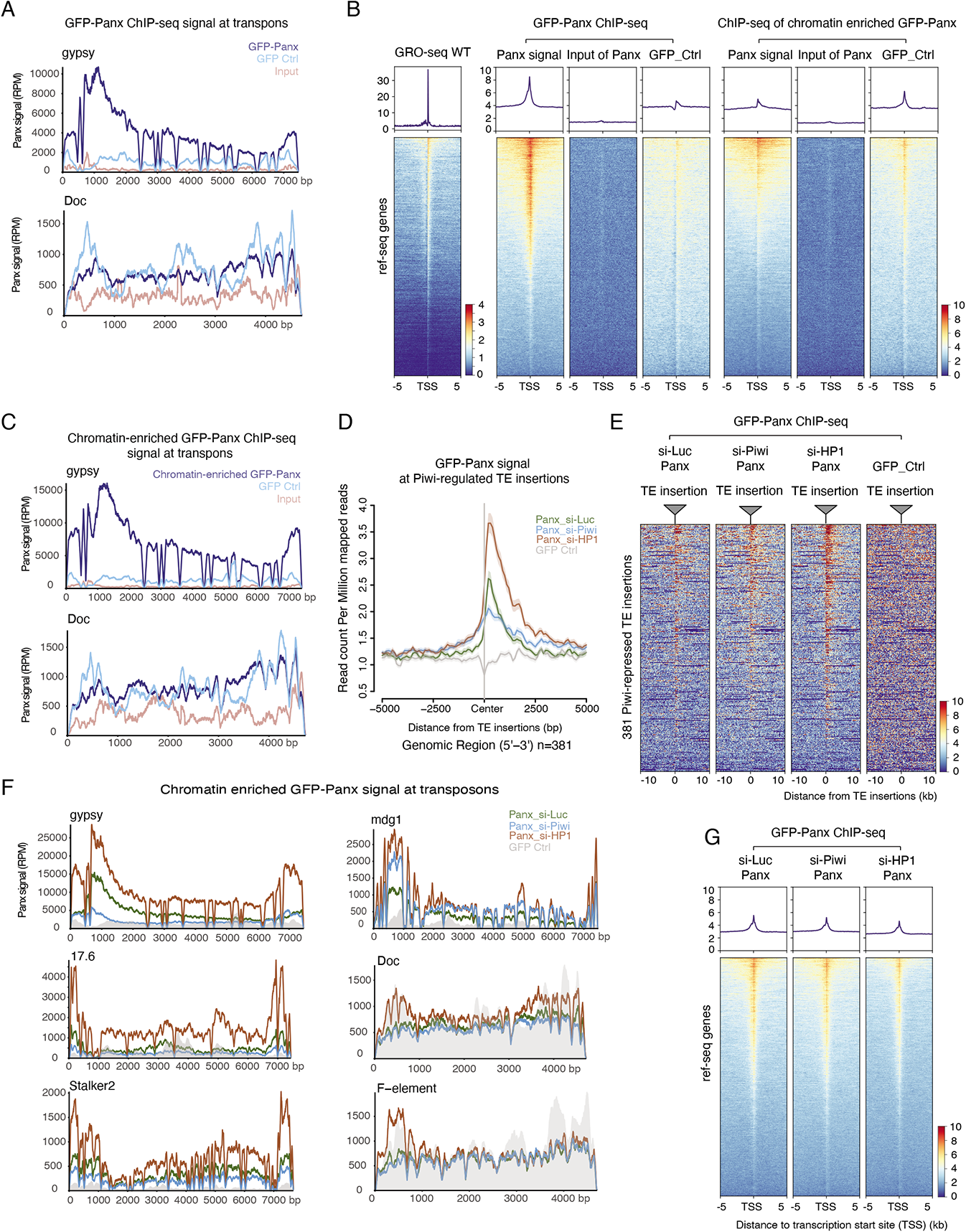
**A**, GFP-Panx ChIP-Seq signal from OSCs expressing GFP-Panx or not (control) at *gypsy* (piRNA targeted) or *Doc* transposons (not targeted). **B**, Heatmap of GRO-seq signal (left), GFP-Panx ChIP-Seq signal (middle) and Chromatin-enriched GFP-Panx ChIP-Seq signal (right) around transcription start sites (TSSs) of expressed genes in OSCs (all heatmaps sorted for maximal GRO-seq signal at the TSS; plots above heatmaps depict the corresponding meta-profiles). **C**, GFP-Panx ChIP-Seq signal from pre-extracted OSCs expressing GFP-Panx or not (control) at *gypsy* (piRNA targeted) or *Doc* transposons (not targeted). **D**, Meta profiles of GFP-Panx ChIP-seq enrichment at genomic regions flanking piRNA-targeted transposon insertions (n = 381 transposon insertions; GFP Ctrl. from a common experiment with Figure 6B). **E**, Heatmap corresponding to meta profile shown in panel D. **F**, GFP-Panx ChIP-Seq signal from pre-extracted OSCs with indicated knockdowns and expressing GFP-Panx or not (control) at indicated piRNA-targeted (*gypsy*, *17.6*, *Stalker2*, *mdg1*) or not targeted transposons (*Doc*, *F*-element). **G**, Heatmap of GFP-Panx ChIP-Seq signal at TSSs of expressed genes in OSCs depleted for the indicated factors.

**Figure S6 (related to Figure 7).**
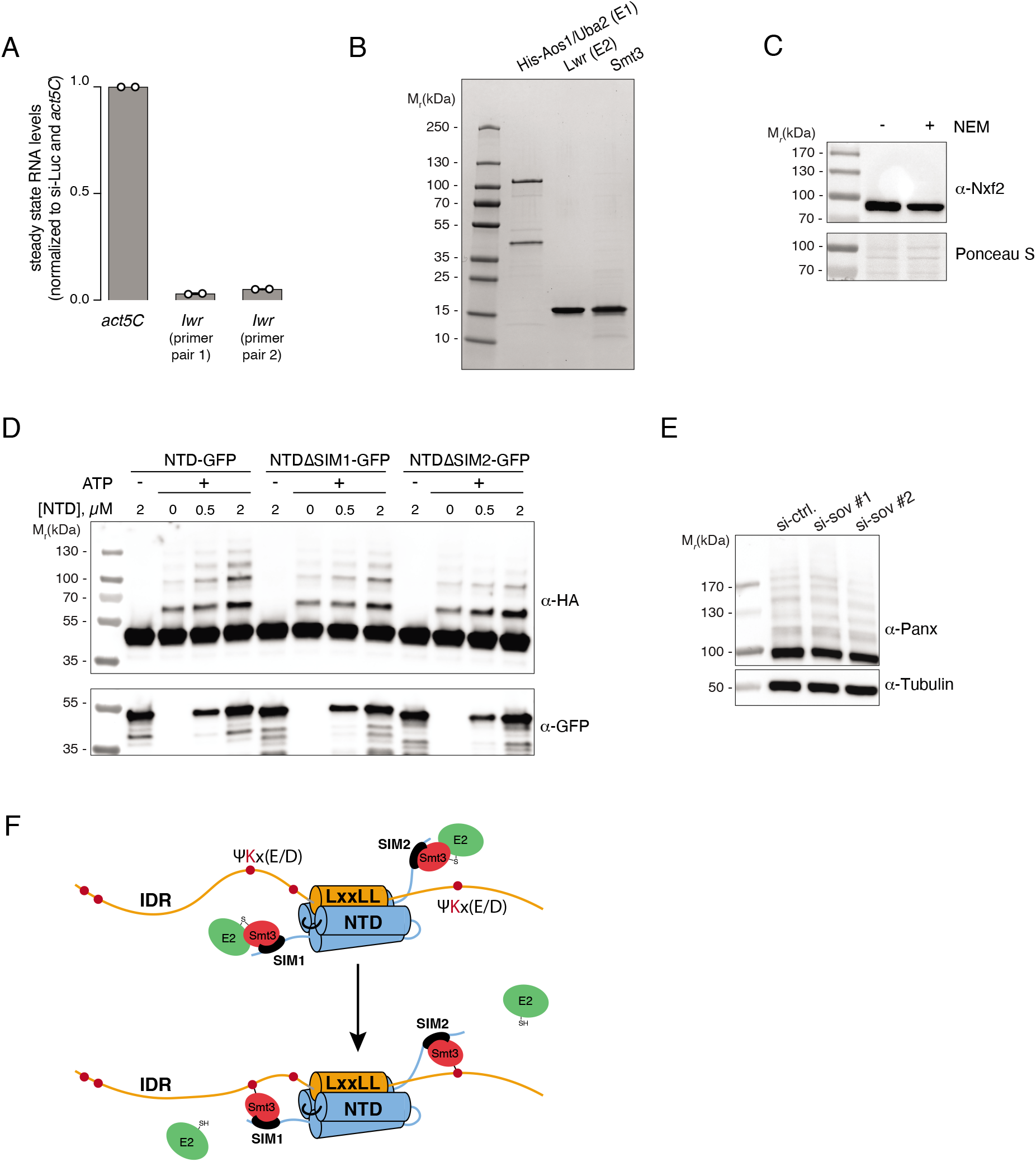
**A,** Transcript levels of *lwr*, measured by RT-qPCR with two amplicons, following siRNA transfection (*n* = 2 biological replicates). **B**, Coomassie-stained SDS-PAGE showing recombinant *Drosophila* Uba2/His6-Aos1, Lwr and Smt3 used in the *in vitro* SUMOylation assays. **C**, Western blot analysis showing endogenous Nxf2 protein in OSC whole cell lysate prepared with or without N-ethylmaleimide (NEM). Staining with Ponceau S serves as loading control. **D**, Western blot analysis showing impact of increasing concentration of Sov NTD variants lacking SIM1 or SIM2 on the efficiency of Panx IDR *in vitro* SUMOylation. **E**, Western blot analysis showing extent of endogenous Panx SUMOylation in OSCs depleted for Sov with two different siRNAs. **F**, Cartoon model of how the Panx–Sov-NTD interaction enhances Panx SUMOylation.

